# Experimental and mathematical modelling of magnetically labelled mesenchymal stromal cell delivery

**DOI:** 10.1101/2020.10.27.356725

**Authors:** E.F. Yeo, H. Markides, A.T. Schade, A.J. Studd, J.M. Oliver, S.L. Waters, A.J. El Haj

## Abstract

A key challenge for stem cell therapies is the delivery of therapeutic cells to the repair site. Magnetic targeting has been proposed as a platform for defining clinical sites of delivery more effectively. In this paper we use a combined *in vitro* experimental and mathematical modelling approach to explore the magnetic targeting of mesenchymal stromal cells (MSCs) labelled with magnetic nanoparticles using an external magnet. This study aims to (i) demonstrate the potential of magnetic tagging for MSC delivery, (ii) examine the effect of red blood cells (RBCs) on MSC capture efficacy and (iii) highlight how mathematical models can provide both insight into mechanics of therapy and predictions about cell targeting *in vivo*.

*In vitro* MSCs are cultured with magnetic nanoparticles and circulated with RBCs over an external magnet. Cell capture efficacy is measured for varying magnetic field strengths and RBC percentages. We use a 2D continuum mathematical model to represent the flow of magnetically tagged MSCs with RBCs. Numerical simulations demonstrate qualitative agreement with experimental results showing better capture with stronger magnetic fields and lower levels of RBCs. We additionally exploit the mathematical model to make hypotheses about the role of extravasation and identify future *in vitro* experiments to quantify this effect.

## Introduction

Mesenchymal stem cells (MSCs) are defined as multipotent, self-renewing cells with the ability to differentiate into several cell lineages. They have been identified as powerful therapeutic agents in the treatment of devastating conditions [3]. To support the swift progression of these therapies to the clinic, research to overcome translational challenges associated with methods of cell delivery is essential. Intravenous (IV) injection is currently the preferred mode of administration for a range of therapies as it is considered to be non-invasive and easy to adopt clinically. However, this delivery method consistently results in the non-specific distribution of injected MSCs throughout the body, most notably in the lungs, liver and spleen, as opposed to the direct delivery of MSCs to the site of injury [22]. Furthermore, pre-clinical and clinical tracking studies have demonstrated only 10% of delivered MSCs persisting at the target sites after IV delivery [22]. To address these challenges, superparamagnetic nanoparticles (MNPs) have been used to control both the delivery to, and retention of, MSCs at the target site [8]. MNPs are internalised by the cells which are then guided to the target site by the application of an external magnet. The magnetic targeting must overcome the forces imposed by the fluid flow and red blood cells (RBCs) which are present within blood vessels.

MNPs are composed of either a magnetite or maghemite core and coated with biocompatible polymers such as dextran or silica [26]. The MNPs used in this study are superparamagnetic, meaning that they are only magnetised in the presence of an external magnetic field. This allows cells with internal MNPs to be controlled close to the external magnet but avoids unwanted magnetic aggregation in other locations. The ability to control the magnetisation of the MNPs means they are now determined as safe by the FDA for use in a similar biomedical application as MRI contrast agents (see [28] and references therein). Additional research has demonstrated that magnetic tagging of MSCs does not adversely affect their viability, proliferation or differentiation potential [27]. The cells retain the magnetic tagging after proliferation, although at a reduced level due to cell division [16] and have been successfully implemented in both *in vitro* [16] and in *in vivo* regenerative studies [28, 38, 40].

Despite the vast potential of magnetic stem cell targeting, a number of challenges hinder its advancement to large scale clinical trials. One such challenge is the effectiveness of magnetic trapping within the physiological environment of a blood vessel which contains a complex mixture of cells including RBCs. At higher levels of RBCs the total viscosity of the blood is higher. This increases the viscous forces on the MSCs from the fluid, potentially reducing magnetic trapping efficacy. Furthermore, *in vitro* studies have shown that magnetic trapping is most effective when the flow is slowest and magnetic field strength is greatest [8, 20]. However, as the magnetic field strength decreases with distance from the magnet, ensuring sufficient magnetic field strength at the target site is a challenge. In addition to the above efficacy challenges, there are also safety challenges which could arise if the MSCs aggregate and obstruct vessels close to the injury site. The interplay between these effects and strategies to avoid or mitigate them must be determined to maintain a safe and effective therapy.

To address these challenges, we use both *in vitro* experimental and mathematical models to examine the effect of RBCs on MSC capture efficacy and to gain insight into minimising vessel obstruction. This offers insights into the interplay between the fluid flow, RBCs, MSCs and magnetic field from a theoretical and practical position. Previous *in vitro* work describes an experimental magnetic trapping model [8] which mimics the *in vivo* vascular network allowing for magnetic platforms to be tested and refined. The system assesses the magnetic trapping efficiency of MNP-labelled MSCs under flow. In this study, we modify this system to include RBCs and assess the effect of varying their concentrations, from 0% to 40% (mimicking physiological conditions), on magnetic MSC capture. The mathematical model enables examination of cellular aggregation on the vessel wall and quantifies how the build up of an aggregate changes the fluid flow and the subsequent MSC capture. This complements *in vitro* work by providing detailed information of the magnetic and fluidic forces on captured cells which are challenging to measure *in vitro*.

The mathematical model represents the *in vitro* system, considering a single 2D channel where MSCs are captured with an external magnet. To examine the capture of large numbers of cells we use a continuum model based on that developed in Grief et al (2005) [14]. We extend the model to include the effect of varying RBC concentration on the MSCs through an effective fluid viscosity [31]. Owing to their size, the cells do not experience Brownian motion. However, collisions between MSCs and RBCs lead to a diffusive type motion known as shear-induced diffusion [23]. This effect was included in Grief et al. (2005). We adapt this model by partitioning the MSCs into two populations. Firstly uncaptured MSCs, which we assume are dilute and therefore do not alter the fluid flow. Secondly captured cells which form a solid mass on the wall of the channel near the magnet. The size of the captured MSC aggregate is determined by its growing boundary. Erosion of the aggregated cells is included proportional to the fluid shear on the aggregate surface.

To access the treatment site, MSCs are required to leave the vessel (extravasate) and migrate towards it. Although the mechanism and kinetics of MSC extravasation are not fully known, experimental measurements of extravasation rates have recorded approximately exponential decay of a fixed amount of cells [21, 24] and linear decrease in cell number over time [4, 21, 24]. Motivated by these observations we test two simple functional forms for the extravasation rate. First, we consider the extravasation rate to be proportional to the number of aggregated cells on the channel wall, and second we consider a constant extravasation velocity. Incorporating extravasation allows the mathematical model to make predictions about its effect on the aggregate size and about the translation of magnetic delivery to *in vivo* scenarios.

This paper is organised as follows. In section 1.1 we present the MSC tagging protocol, the *in vitro* flow system and methods for subsequent analysis of trapped cells. In section 1.2 we then explain in brief the mathematical model and numerical method for solving it, with the full details given in the Appendix. In section 2 we present and compare the results of the *in vitro* trapping, numerical simulations of the mathematical model and predictive results for the effect of extravasation on MSC capture. We conclude with a discussion of both models, their successes and limitations. We also suggest *in vitro* experiments which could be used to validate and extend these models (sections 3 and 4).

## 1 Materials and Methods

### 1.1 Experimental *In Vitro* Model

#### Expansion and MNP labelling of oMSCs (ovine Mesenchymal Stem Cells)

Bone marrow derived oMSCs (P3-5) were used throughout this study. Cells were cultured in expansion media consisting of Dulbecco’s modified eagle medium (DMEM) supplemented with 10% fetal bovine serum (FBS), 1% L-glutamine and 1% penicillin-streptomycin and incubated at 37°C with 5% *CO*_2_ until 90% confluent. To label cells with SiMAG (Chemicell, Germany), a commercially available 1000nm, Silanol coated MNP, cells were initially plated at a seeding density of 1.4 × 10^4^ cells per cm^2^ and allowed to attach overnight as described by Markides and Harrison et al. [16]. Labelling solution was then prepared by re-suspending SiMAG particles (1000 nm, Chemicell, Germany) in serum free media (SFM) to achieve a final iron concentration of 10*μ*g [Fe] per 2 × 10^5^ cells and allowed to incubate overnight to encourage passive uptake. Following internalisation, cells were washed three times with phosphate buffered solution (PBS) to remove unbound MNPs and prepared for trapping studies.

#### Prussian Blue detection of MNPs

Prussian blue is an iron-based stain routinely used to identify the presence of MNPs. SiMAG-labelled cells were stained with Prussian blue to validate the internalisation of SiMAG by oMSCs. Here, labelled cells were fixed with formalin (10min; RT) then treated with a 1:1 solution of 10% aqueous solution of potassium hexacyanoferrate and a 20% aqueous solution of concentrated Hydrochloric acid (HCL) (20 min; RM). The presence of MNPs is observed by bright blue staining when imaged by bright-field microscopy (EVOS XL Core Cell Imaging System).

#### *In vitro* experimental model

To experimentally evaluate the effect of a) magnet strength and b) RBC concentration on magnetic mediated cell trapping, an *in vitro* magnetic trapping model was modified from El Haj et al. (2015) [8] (Figure 1 A). We extend analysis from [8] through inclusion of RBCs. We then use this system along with a new system with stronger, larger magnets. In brief, the trapping systems consist of neodymium iron boron magnets evenly spread out and embedded across either a polystyrene casing with three 0.4T magnets (Figure 1 B) or a purpose-built plastic box with four 0.2T magnets (Figure 1 A). SiMAG-labelled and unlabelled cells (1 × 10^6^) were then flowed through 30 cm length PVC tubing (I.D. 1.65mm, Tygon® E-3603) via a 5ml reservoir and tubes secured over each magnet. Each set of tubes were attached to a digital peristaltic pump (Ismatec) set at a pump rate of 1ml/min for 30 min.

**Figure 1:**
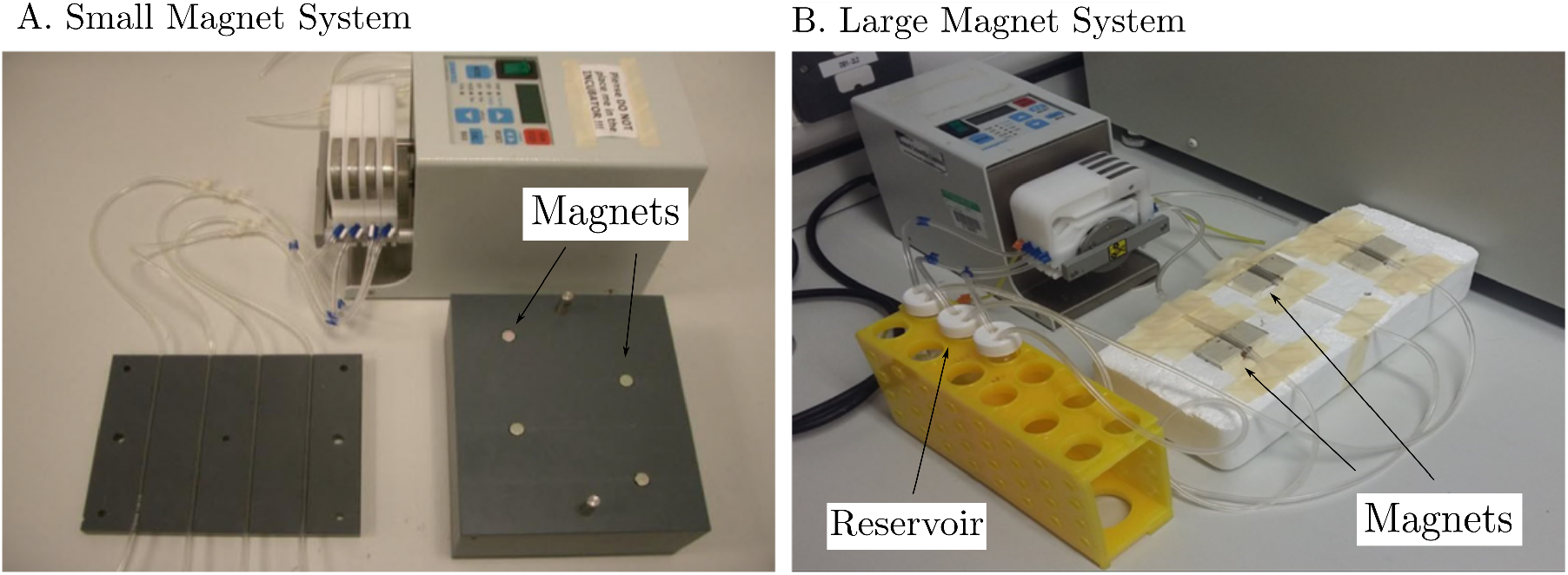
*In vitro* magnetic trapping systems. A) This represents the “small magnet” system, El Haj et al. (2015) whilst B) represents the “large magnet” system.

SiMAG labelled or unlabelled cells were re-suspended in the desired RBC concentration (0, 5, 10, 20 or 40%) using sheep blood in Alsever’s (TCS biosciences, 20% stock concentration) prior to running samples through the system. To achieve 40% RBC concentrations, the stock blood was centrifuged for 30 minutes at 1000 rpm, and half the supernatant removed prior to re-suspension. In addition, to prevent the blood from clotting, a heparin solution was prepared and 0.2 ml added to the cell suspension. To reduce any trapping which may occur due to the natural adhesion of cells to the tubing walls, heparin was further pumped through the system to line all tubes for 5 min prior to sample addition. Upon completion, 1 cm sections of tubing were taken from each edge of the magnet (where the magnetic field is strongest) and a complimentary 1 cm control section of tubing taken from a distance sufficiently far as not to be affected by the magnets. Un-trapped cells remaining in the reservoir were further processed for analysis.

#### RBC lysis

To accurately quantify the percentage of MSCs trapped, it was necessary to remove all the RBCs. This was achieved using Red Blood Lysis buffer (Sigma-Aldrich, hybrid-max™) which preferentially lyses RBCs from a mixed population of cells. Cells were treated with ice-cold Red Blood Lysis buffer for 3-5 min with gentle agitation or until an obvious shift in colour from bright red to brown was observed (demonstrating RBC oxidation and successful lysis). The cells were then centrifuged for 5 min at 1200 rpm. This was repeated until all RBCs were removed and lysed. Samples were finally stored in 100*μ*L of 0.1% triton-x at −80°C, and freeze-thawed three times before analysing by Picogreen.

#### Quantification of cell aggregate

The PicoGreen (Quanti-iT™ PicogreenQ® dsDNA Kit (Invitrogen)) assay was used to quantity double stranded DNA (dsDNA) present in a sample as recommended by the manufacturer. This is a good indication of cell number and used to determine the proportion of trapped cells relative to total cell number.

### 1.2 Mathematical model

We consider cell delivery in a single straight vessel which we approximate by a two-dimensional channel of height 2*d* and length 2*l* containing fluid, MSCs and RBCs. We distinguish between MSCs in the flow, modelled by a concentration, and MSCs trapped in an aggregate on the vessel wall, which we model as a solid. We assume the cells in the channel are sufficiently dilute that they do not alter the fluid flow. The extent of the aggregate is defined by a moving boundary, the evolution of which is determined by the MSC concentration and fluid flow in the channel. This setup is shown in Figure 2. Full details of the mathematical model can be found in the Appendix.

**Figure 2:**
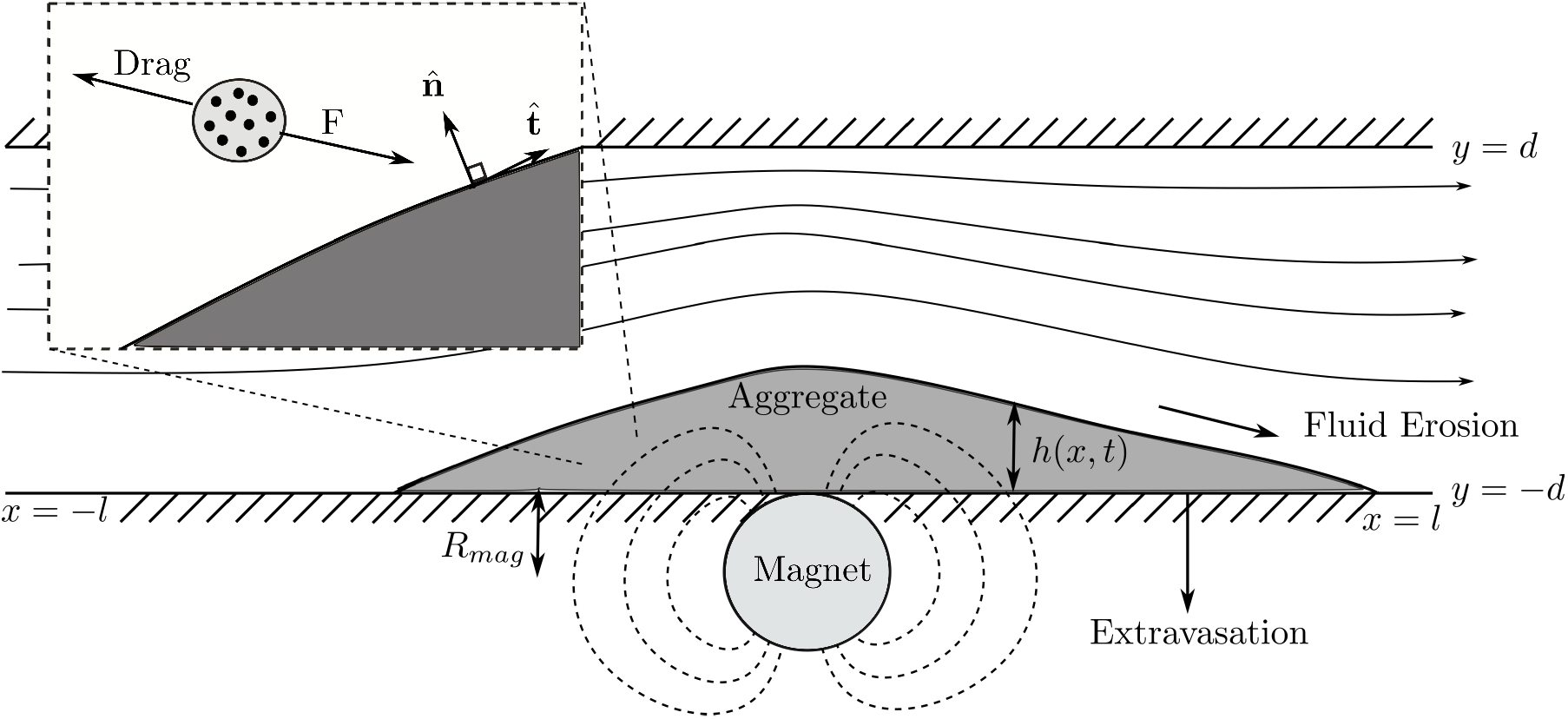
The mathematical model setup: a channel with length 2*l* and height 2*d* is located above a magnet of radius *R*_mag_, we use Cartesian coordinates (*x, y*). The boundary of the aggregate lies at *y* = −*d* + *h*(*x, t*). Illustrative fluid streamlines and magnetic field lines (dashed) are shown. Inset shows a cell tagged with NNPs travelling under the action of magnetic force ***F*** and fluid drag.

#### Fluid flow

We neglect fluid inertial effects since the therapy aims to capture MSCs in arterioles (Reynolds number 0.03 [14]) prior to their extravasation. Hence we model the fluid flow with the Stokes and continuity equations

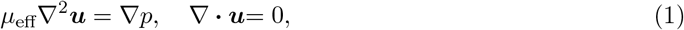

where ***u*** and *p* are the fluid velocity and pressure respectively and *μ*_eff_ is an effective viscosity.

The viscosity of blood depends nonlinearly on RBC volume fraction and the diameter of the vessel. The haematocrit, H is defined as the average volume fraction of RBC in the cross-section of the pipe. We use an empirical fitting for the observed viscosity of a suspension of RBC for varying haematocrit *in vitro* [31, 32], shown in Figure 3a. This includes the average viscosity change from RBC marginalisation as vessel radius increases and further viscosity increase as *H* increases.

**Figure 3:**
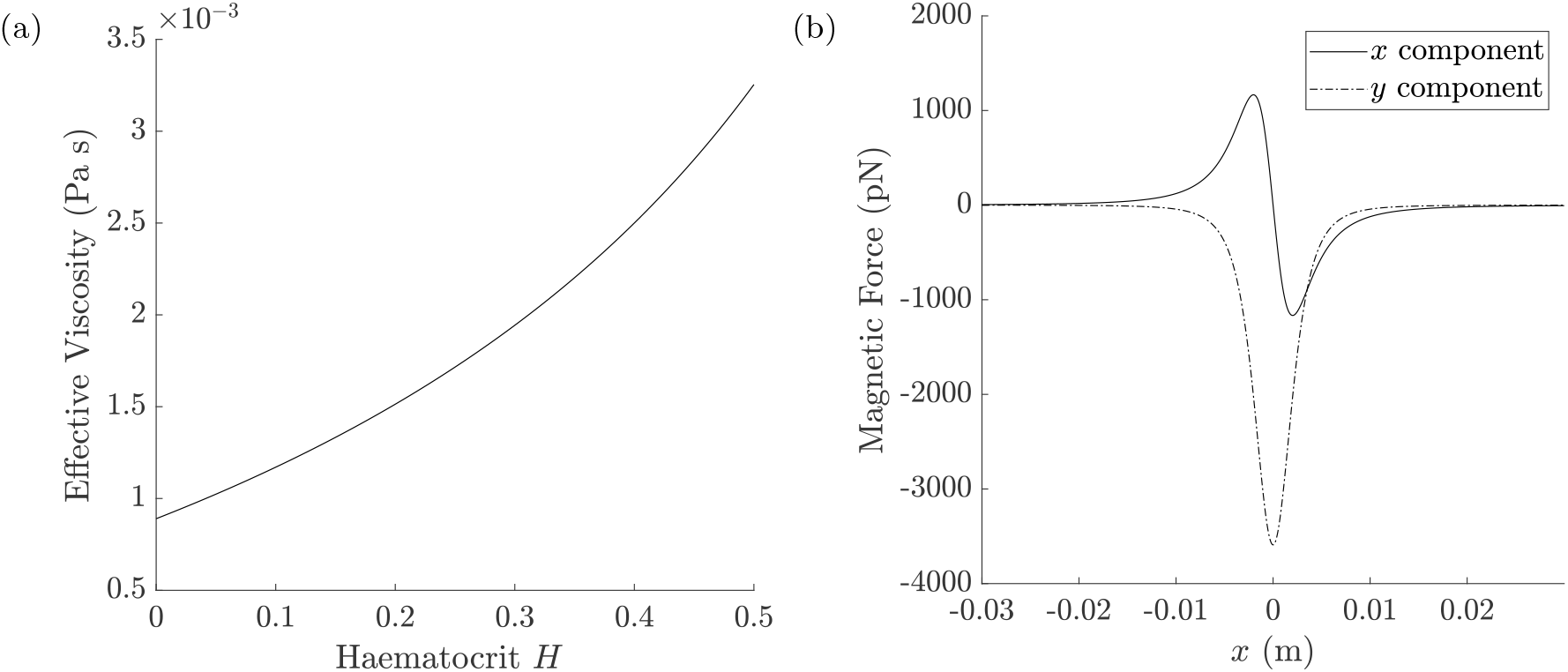
(a) Fitting of viscosity of whole blood *in vitro* in a vessel radius 825*μ*m for varying haematocrit [31], definition in Appendix, Eq. (6). (b) Magnetic force on an MSC at *y* = 0 with 30 implanted MNPS from a magnet with radius *R*_mag_ = 0.35*cm* with *B*_*max*_ = 0.4T as in experimental setup, definition in Appendix, Eq. (11).

The boundary conditions for the flow are no-slip and no normal flow on the stationary, impermeable, channel walls. The aggregate-fluid interface, the fluid moves with the velocity of the interface, 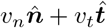 denoted. The vectors 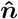 and 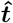 are the inward (relative to the fluid) unit normal and unit tangent vectors, see Figure 2. The height of the cell aggregate *h* is determined via the kinematic boundary condition. Since the aggregate is modelled as solid, no-slip also applies on the aggregate boundary. The flow is driven by a unidirectional, parabolic flow with maximum *u*\* at the inlet. At the outlet, we prescribe atmospheric pressure and no tangential stress.

#### Magnetic force

The force ***F*** on a single MNP in an external field ***B*** is ***F*** = ***m*** · ∇***B***. Here ***m***(***B***) is the magnetic moment of the MNP which defines its strength and alignment with the external magnetic field, and is a function of the magnetic field.

In this model, we neglect interactions between the MNPs. Furthermore, since the maximum field strength used *in vitro* is large compared to the required saturation strength of the MNPs (estimated as 0.05T), we assume the nanoparticles are permanently saturated. Under these assumptions, the moment of a single MNP with volume *V_p_* is 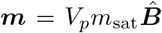 where *m*_sat_ is the saturation value of the MNP and 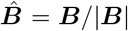 is the normalised magnetic field. Owing to the small size of the cell we model the magnetic moment of a cell tagged with *n* MNPs as a linear multiple of the moment for an individual MNP as in [12]. Hence the force on a cell is defined 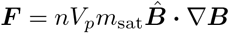.

The external magnetic field is generated by an infinite cylindrical magnet of radius *R*_mag_ located directly below the channel, oriented perpendicular to it and with magnetisation *M* in the *y* direction. This allows the magnetic field (and also the force on a cell) to be expressed analytically [11], see appendix Eq. (11). The magnetic force experienced by a tagged cell is shown in Figure 3b.

#### Cell transport

The MSC volume concentration *c* is governed by an advection-diffusion equation, such that the cells move with the fluid velocity ***u*** and an induced magnetic velocity ***u***_*m*_. This is defined by balancing the magnetic force from the external magnet with viscous drag from the surrounding fluid giving ***u***_*m*_ = ***F*** /6*πμ*_eff_*r* where *r* is a typical MSC radius. We include shear induced diffusion proportional to 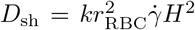 [23] where 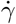 is the local shear rate defined as the 2-norm of the strain rate tensor, *r*_RBC_ is the radius of a typical RBC, and *k* = 0.5 is a constant (obtained through asymptotic analysis at low RBC volume fraction [23]). This leads to the following advection-diffusion equation from Grief et al. (2005) [14] modified to remove Brownian motion

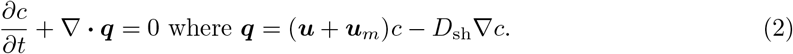

Cells arrive at the inlet between *t* = 3*t*\* and *t* = 35*t*\* with concentration *c*_in_(*t*), where *t*\* = 0.11s is a typical timescale. The concentration *c*_in_(*t*) is defined by *c*_in_ = *C_in_*(tanh (*t* − 3*t*\*) −tanh (*t* − 35*t*\*))*/*2 where *C*_*in*_ is a constant. Initially we take *c* = 0 throughout the channel. We impose zero total flux on the top of the channel and zero diffusive flux at the channel exit, allowing cells to exit the channel with the flow. On the base of the channel the flux *J* into the aggregate relative to the moving boundary is taken as

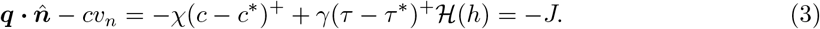

The first term represents uptake at a speed *χ* into the aggregate above a critical concentration threshold *c*\*. Shear-stress erosion is modelled by the second term. It is taken proportional to the difference of the shear stress 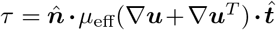 from the erosion threshold *τ* *. We specify the positive part to ensure that erosion is never negative, and the Heaviside function 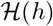 ensures the boundary does not grow downwards when there is no aggregate present. The parameter *γ* models the sensitivity of the aggregate to shear stress erosion.

#### Aggregate growth

The aggregate interface grows as a result of the arriving cell flux. We note that the cell flux can be negative, if fluid shear is high enough, corresponding to cells re-entering the channel from the aggregate. We assume that the aggregate grows solely in the direction normal to its boundary with velocity *υ_n_*, such that the tangential velocity is zero. The height is reduced by extravasation of cells through the channel wall. The aggregate growth velocity is hence

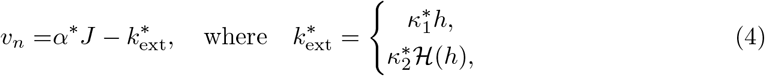

where *J* is the flux of cells into the aggregate, the coefficient *α* is the growth velocity scale of the aggregate proportional to the arrival of cells and *k*_ext_ is the extravasation function. We consider two forms of extravasation: firstly aggregation proportional to the height of cells on the base, marked with (∗) and constant extravasation marked with (∗∗). The constant *κ*_1_ determines the rate at which a collection of cells extravasate, whereas *κ*_2_ corresponds to the speed at which a single cell crosses the vessel wall. Initially we take the aggregate height to be zero.

#### Parameterisation

Parameters for the model are shown in Table 1. As far as possible parameters have been directly matched to *in vitro* experiments. Remaining parameters have been estimated. Aggregate parameters, shown in Table 2, are unknown and quantification of these would require further experimentation. Our strategy here is to fix *α, χ, c*\* and *τ*\* at values which lead to physically realistic solutions. This is defined by solutions where the aggregate growth speed is less than the fluid speed, the cells are able to withstand some shear stress and the cells build up before a solid aggregate forms. We then vary *κ*_1_, *κ*_2_ and *γ* to demonstrate the qualitatively different behaviours.

**Table 1:** Physical parameters and units. Those with * are taken directly from *in vitro* setup whereas those marked with ^†^ have been estimated. To consider the relevant section of the system where MSCs are captured the half length of the channel is *l* = 1.65cm.

| Parameter | Definition | Units | Value |
| --- | --- | --- | --- |
| $C_{\text{in}}$ | Inlet concentration (volume concentration) | - | $8.4 \times 10^{-4*}$ |
| $l$ | Channel half length | m | $1.65 \times 10^{-2\dagger}$ |
| $d$ | Channel half height | m | $8.25 \times 10^{-4*}$ |
| $H$ | Haematocrit | - | $0 - 0.4^*$ |
| $n$ | Number of MNPs in cell | - | $30^\dagger$ |
| $\mu_{\text{water}}$ | Viscosity of water | Pa s | $8.9 \times 10^{-3}$ [2] |
| $R_{\text{mag}}$ | Magnet radius | m | $3.5 \times 10^{-4\dagger}$ |
| $r_{\text{RBC}}$ | RBC radius | m | $3.6 \times 10^{-6}$ [37] |
| $r$ | MSC radius | m | $1 \times 10^{-5*}$ |
| $u^*$ | Maximum inlet velocity | m/s | $0.0078^*$ |
| $\mu_0$ | Permeability of vacuum | N/A<sup>2</sup> | $4\pi \times 10^{-7}$ [12] |
| $V_p$ | Volume of NP | m<sup>3</sup> | $4.18 \times 10^{-18*}$ |
| $M$ | Magnetic saturation of external magnet | A/m | $[1.6, 3.2, 4.8, 6.7] \times 10^{5\dagger}$ |
| $m_{\text{sat}}$ | Magnetic saturation of NP | A/m | $1.25 \times 10^{5\dagger}$ |
| $k_B$ | Boltzmann constant | J/K | $1.38 \times 10^{-28}$ |
| $T$ | Temperature | K | 295 |
| $k$ | Shear induced diffusion coefficient | - | 0.5 [23] |

**Table 2:**
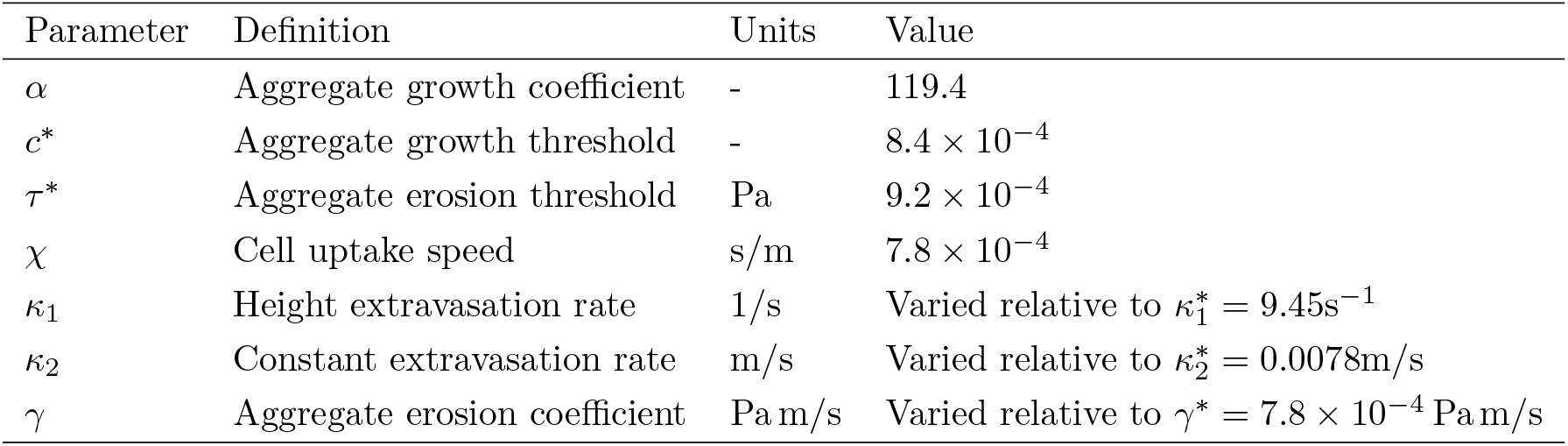
Aggregate model parameters with units. We fix some parameters to achieve physically realistic solutions and vary *κ*_1_*, κ*_2_ and *γ* to reveal the solution behaviour. These are varied relative to typical values marked with an asterisk. We note that *α* is dimensionless since we consider volume concentration of MSCs.

| Parameter | Definition | Units | Value |
| --- | --- | --- | --- |
| $\alpha$ | Aggregate growth coefficient | - | 119.4 |
| $c^*$ | Aggregate growth threshold | - | $8.4 \times 10^{-4}$ |
| $\tau^*$ | Aggregate erosion threshold | Pa | $9.2 \times 10^{-4}$ |
| $\chi$ | Cell uptake speed | s/m | $7.8 \times 10^{-4}$ |
| $\kappa_1$ | Height extravasation rate | 1/s | Varied relative to $\kappa_1^* = 9.45\text{s}^{-1}$ |
| $\kappa_2$ | Constant extravasation rate | m/s | Varied relative to $\kappa_2^* = 0.0078\text{m/s}$ |
| $\gamma$ | Aggregate erosion coefficient | Pa m/s | Varied relative to $\gamma^* = 7.8 \times 10^{-4} \text{Pa m/s}$ |

#### Numerical solution

To solve the governing equations we used finite element formulation of the PDEs coupled to the moving lower boundary through the use of Arbitrary Lagrangian-Eulerian Method (ALE) [18]. All simulations were carried out using a commercial software COMSOL [6]. Taylor-Hood elements (quadratic and linear Lagrange) were used for the fluid velocity and pressure owing to their established stability properties [36]. The advection-diffusion equation was solved with quadratic elements. We use inbuilt adaptive timestepping method IDA. This uses implicit backwards differentiation formula of order 1 or 2 depending on local error. We set the relative tolerance to 0.005 and the absolute tolerance to 0.05 scaled on the concentration solution, see details in COMSOL documentation [6].

Inclusion of shear-induced diffusion is a significant challenge numerically as the shear rate vanishes in the centre of the channel. This renders the cell transport equation hyperbolic and the finite element method unstable which leads to spurious oscillations in the numerical solution. To ensure the simulations are tractable, the shear-induced diffusion is approximated by larger isotropic diffusion, removing the dependence of the diffusion coefficient on fluid shear and haematocrit.

The system was solved on an evolving domain using an unstructured quadrilateral mesh. We allowed free movement of mesh points throughout the entire domain and did not require remeshing. All Heaviside functions which are approximated by a fifth-order polynomial. Sensitivity tests and mesh convergence tests can be found in the Appendix.

## 2 Results

### Mathematical model application to *in vitro* delivery: numerical results demonstrate key stages of cell capture

We first consider the mathematical model in the context of the *in vitro* experiments such that there is no extravasation. The transport of MSCs and the evolution of the cell aggregate on the wall, as predicted by the model, are shown in Fig. 4. First, shown in Fig. 4a, the cells are advected with the flow. Then, once the cells are within range of the magnet, some of the cells are captured on the base of the channel. As more cells arrive the MSCs form an aggregate (Fig. 4b). Note that since the magnetic field is strong the aggregate forms upstream of the centre of the magnet (*x* = 0). As more cells arrive the aggregate significantly obstructs the channel (Fig. 4c). To preserve the flux through the channel the flow speeds up over the peak and increases shear stress on the aggregate. Additionally, as the source of cells at the inlet ends the aggregate decreases in size as the cells are both eroded from the surface by shear-stress and extravasate via height dependent extravasation (Fig. 4d). Once the aggregate is smaller it translates upstream to centre at the location of the magnet. Since extravasation is present the remaining cells leave the channel through the wall.

**Figure 4:**
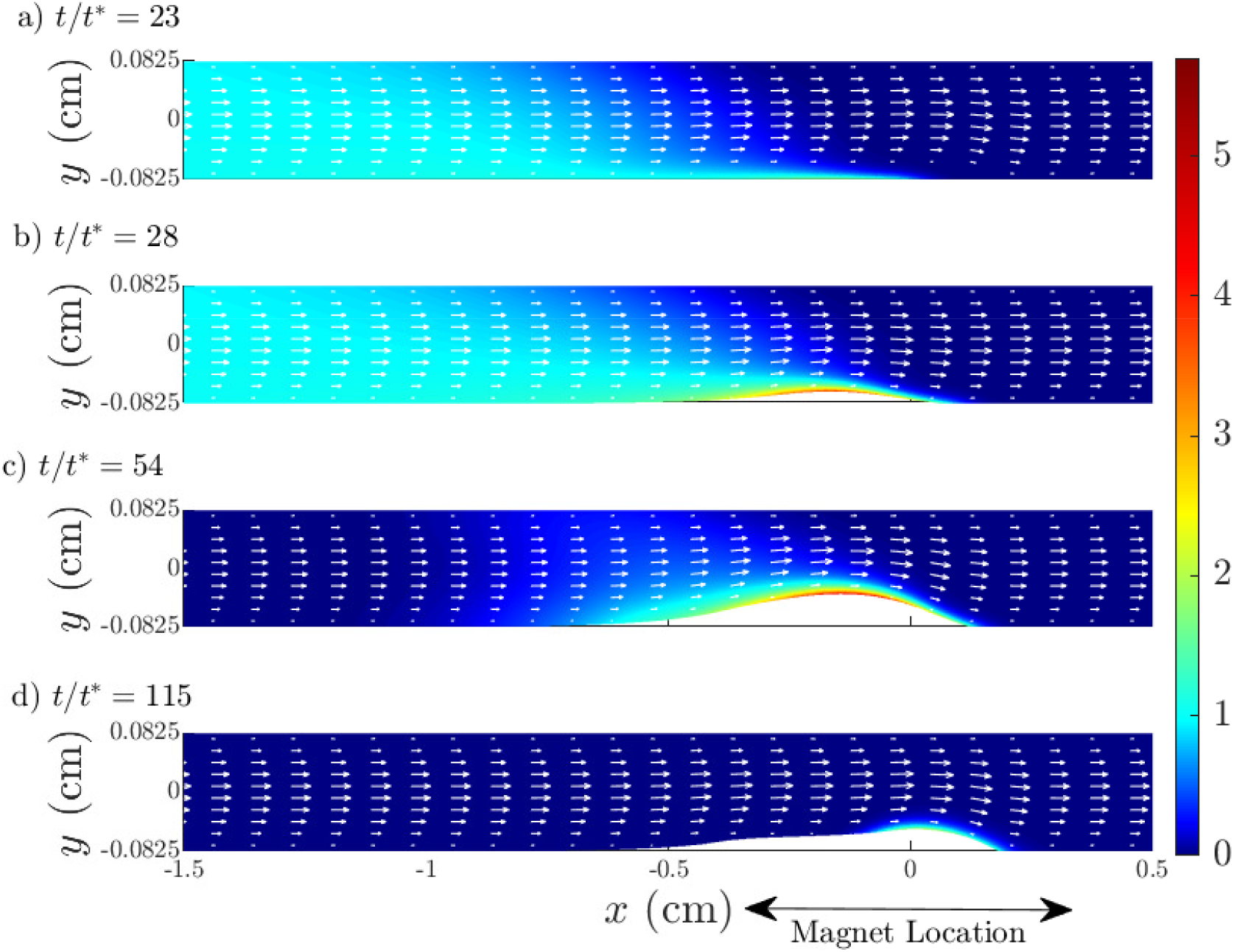
Transport of MSCs and cell aggregation at the channel wall is shown in snapshots of cell concentration relative to the inlet concentration together with fluid velocity field. Section of channel occupying 2cm over the magnet is shown. (a) MSCs arrive via fluid advection then begin to build up on the base close to the magnet, *t/t*\* = 23; (b) MSC level increases (shown by colour bar) and solid aggregate begins to form at *t/t*\* = 33; (c) the aggregate extent grows and the channel is obstructed significantly at *t/t*\* = 54; (d) once the inlet source of MSCs ends, the captured MSCs are eroded by fluid shear and cells extravasate (via height dependent extravasation) out of the channel (both shown by decreasing aggregate size) by *t/t*\* = 115. Parameter values: *H* = 0.2, *B_max_* = 0.4*T, κ*_1_ = 0.01*κ*\**, γ* = 0.1*γ*\*, *t*\* = 0.11s all other parameters as in Table 2.

The stages are illustrated clearly in Fig. 5 illustrating the velocity of the aggregate boundary at *x* = −0.17cm. The relative size of the growth rate compared to erosion and extravasation rates determines the capture efficacy of MSCs and whether the channel will potentially block.

**Figure 5:**
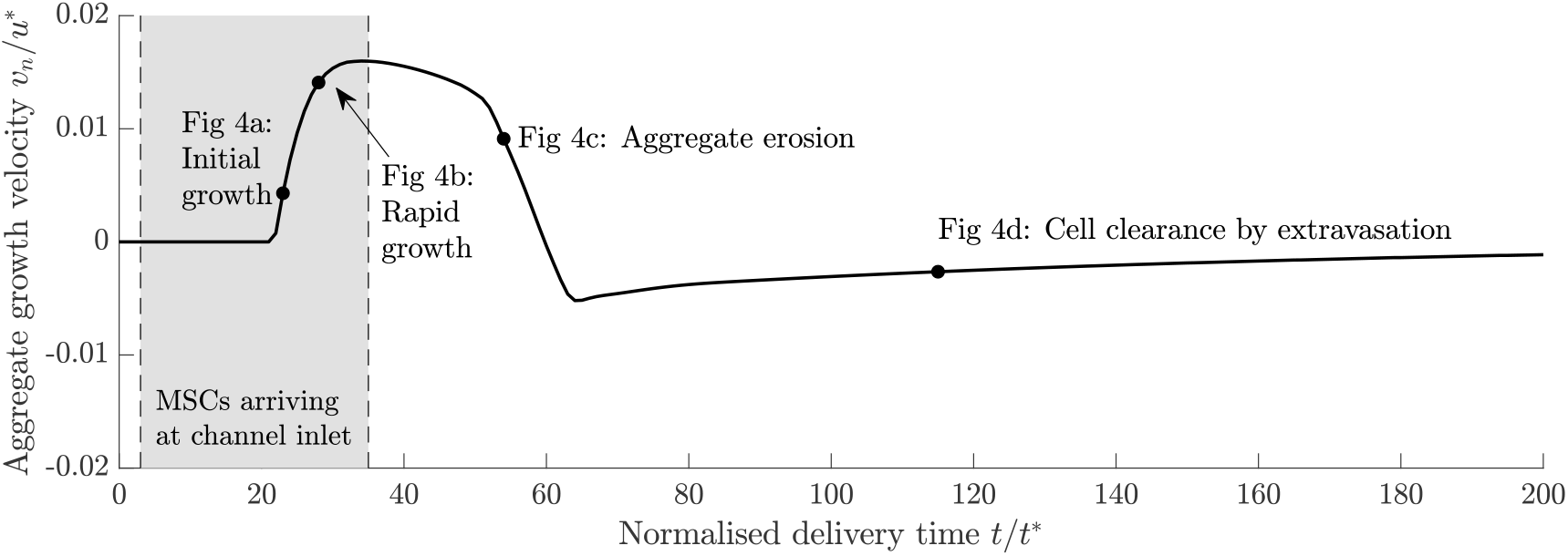
Aggregate interface velocity at *x* = *−*0.17cm as MSCs are captured. Negative velocity indicates when the aggregate is shrinking. The points correspond to snapshots of the system in Fig. 4. Shaded region illustrates when MSCs are arriving at the channel inlet. Parameter values as in Fig. 4.

### SiMAG is efficiently taken up by o-MSCs and a minimum labelling efficiency is required to promote capture of cells on the vessel wall

Prussian blue staining successfully confirmed the uptake of SiMAG by oMSCs as seen by the blue staining in the internal regions of the cells (Fig. 6B) when compared to unlabelled cells (Fig. 6A). Analysis of the cells which were captured on the surface by the magnet revealed significantly more blue staining in the captured fractions (Fig. 6D) compared to their uncaptured counterparts (Fig. 6C). This implies that a minimum particle uptake is required for efficient capture of the cells to the vessel wall. This is estimated to be 15pg/cell based on previous unpublished work.

**Figure 6:**
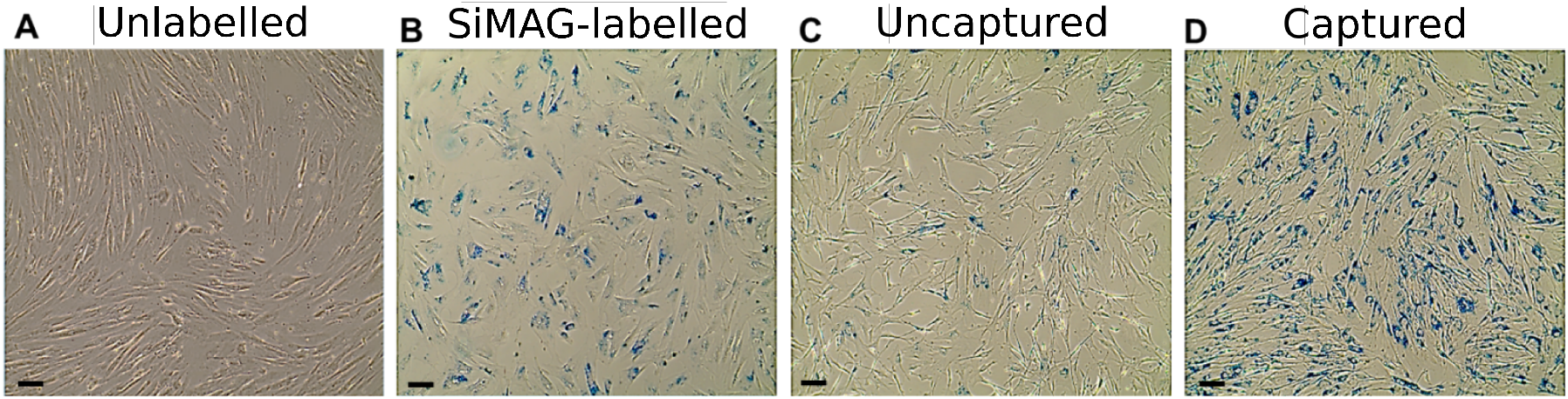
Assessment of SiMAG internalisation by Prussian blue staining which stains for the iron oxide core of the MNP. Prussian blue staining of A) oMSCs incubated without SiMAG (unlabelled). B) oMSCs incubated with 10*μ*g/ml of SiMAG in serum free media overnight. C) oMSCs which have passed through the magnetic trapping system in 0% blood for 30min but were not trapped. D) oMSCs which were trapped by the magnetic trapping system in 0% blood after 30 min. Scale bar = 100*μ*m.

### High levels of RBCs limits the efficiency of magnetic trapping of MNP-labelled oMSCs

SiMAG labelled oMSCs are efficiently trapped by the external magnet when compared to unlabelled control cells (Fig. 7). This is most efficient when cells are passed through in media alone (no RBCs) irrespective of magnet size with approximately 50% of cells trapped at the magnet site. The trapping efficiency of the system significantly drops in the presence of RBCs but to a greater extent in the smaller magnet than the larger magnet. A further 50% drop in trapping efficiency is determined with RBC concentrations ranging from 5-20% compared to 1% in the larger magnet system (Fig. 7B). A degree of trapping is detected in unlabelled control cells at the magnet site with increasing RBC concentration, this is likely to be attributed to increased stickiness from the RBCs.

**Figure 7:**
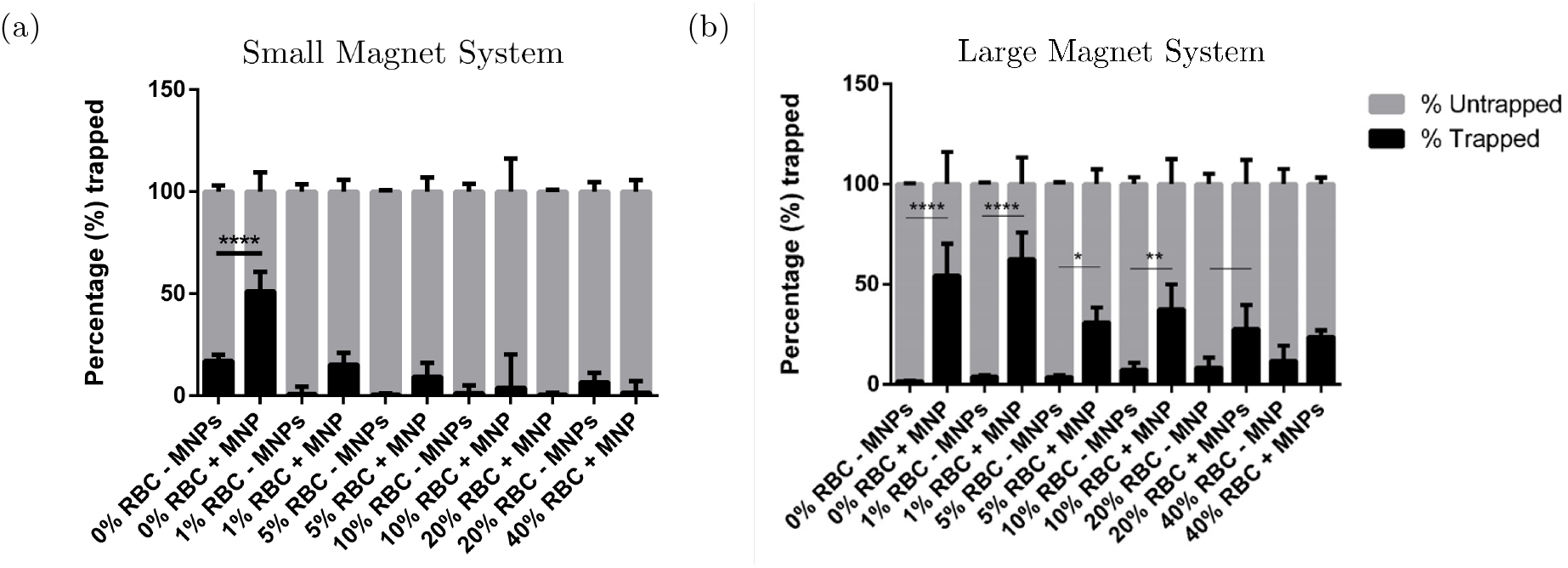
Graphs demonstrating the percentage of SiMAG labelled or unlabelled oMSCs trapped by either A) the small magnet (n=4) or B) the large magnet system with increasing RBC concentrations (0, 1, 5, 20 and 40%) after 30 min of flow (n=3). Significance is determined by two-way Anova where **** is p <0.0001, *** is p<0.001, ** is p<0.01 and * is p<0.05.

### Stronger magnetic fields capture more cells further upstream

For a given flux in the channel, the mathematical model demonstrates increasing the magnetic field strength increases cell capture, as shown in Figure 8a by an increase in both the height and the extent of the aggregate. Furthermore, for weaker magnetic fields cells enter the aggregate upstream while other cells flow over the aggregate and are captured downstream, thus ensuring approximately symmetric capture of cells around the magnet. However, as the magnetic field strength increases the cells no longer flow over the aggregate, all entering upstream of the peak and the downstream slope becomes steeper.

**Figure 8:**
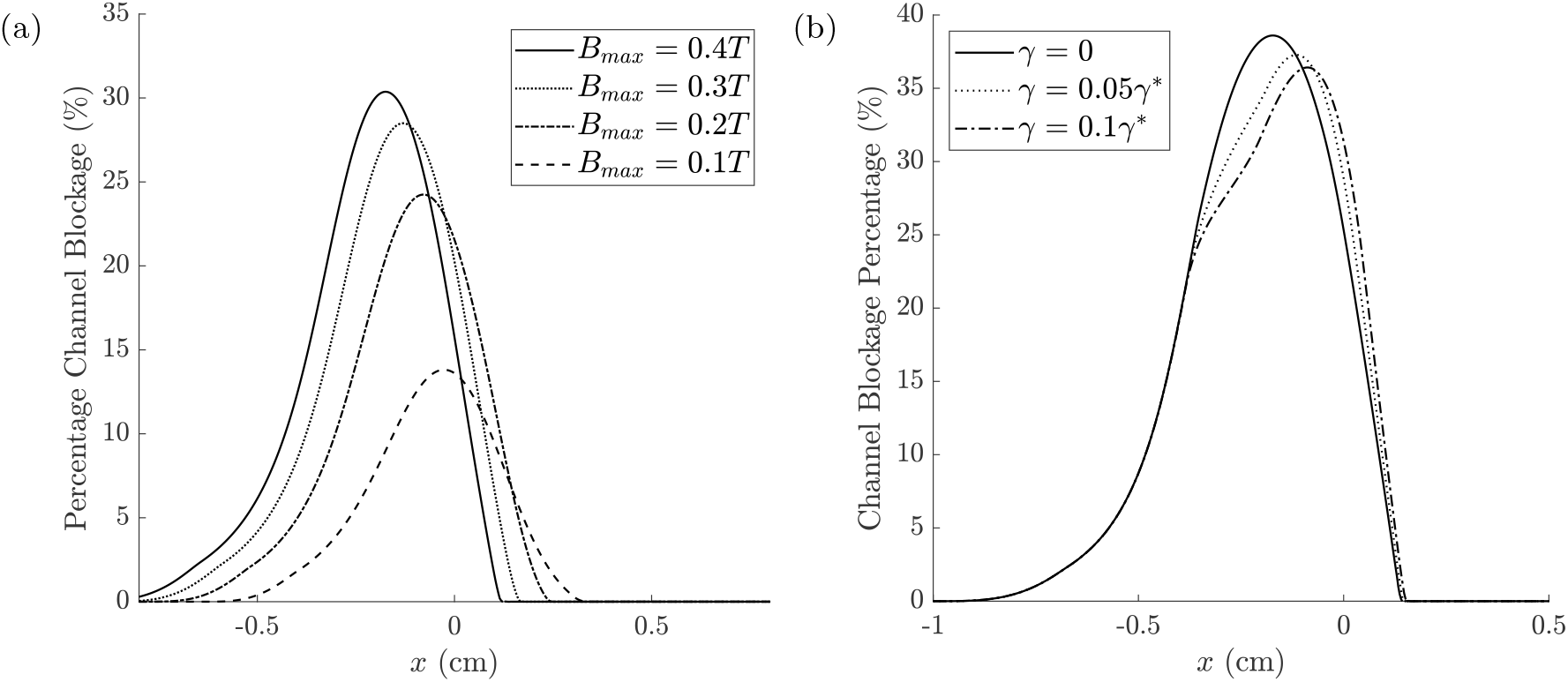
(a) Aggregate interface profiles increase for increasing magnetic field strength. Parameter values: *H* = 0.2, *κ*_1_ = 0 (no extravasation), *γ* = 0 (no erosion) at *t* = 50*t*\* where *t*\* = 0.11s. (b) Aggregate peak decreases for increasing erosion parameter. Parameter values: *H* = 0.2, *B_max_* = 0.4*T, κ*_1_ = 0 (no extravasation), and *t* = 80*t*\* where *t*\* = 0.11s all others as in Table 2.

### Increased cell susceptibility to fluid erosion leads to less cell capture

For larger values of the susceptibility parameter *γ* the aggregate peak, where the fluid shear is greatest, is reduced. This forces aggregated cells to migrate downstream, where the magnetic field is strong enough to overcome erosion. The threshold of erosion which cells are assumed to withstand is set at 10% higher than the fluid shear on the walls in the absence of an aggregate. Once the aggregate exceeds 25% of the channel height erosion begins to remove the cells arriving on the upstream slope.

### Mathematical model recreates *in vitro* trapping reduction with haematocrit

In Fig. 9a we compare the maximum channel blockage produced for varying haematocrit levels at the magnetic field strengths used *in vitro*. For both strengths of magnetic fields, increasing haematocrit decreases the maximum aggregate height captured, reflecting the results obtain *in vitro* (Fig. 7). Increasing haematocrit levels increases the viscosity of the fluid which in turn decreases the relative effect of the magnetic force, as the drag on the cells increases. It additionally increases the strength of erosion on the aggregate. For both magnetic field strengths, the maximum blockage possible in the range of 0%-40% RBCs is 40%. Since fluid erosion increases as the channel is increasingly obstructed, new arriving cells are not able to withstand the adverse flow effects and do not attach to the aggregate. This saturation of cell capture is also clear *in vitro* although this effect occurs at a quantitatively different concentration of RBCs.

**Figure 9:**
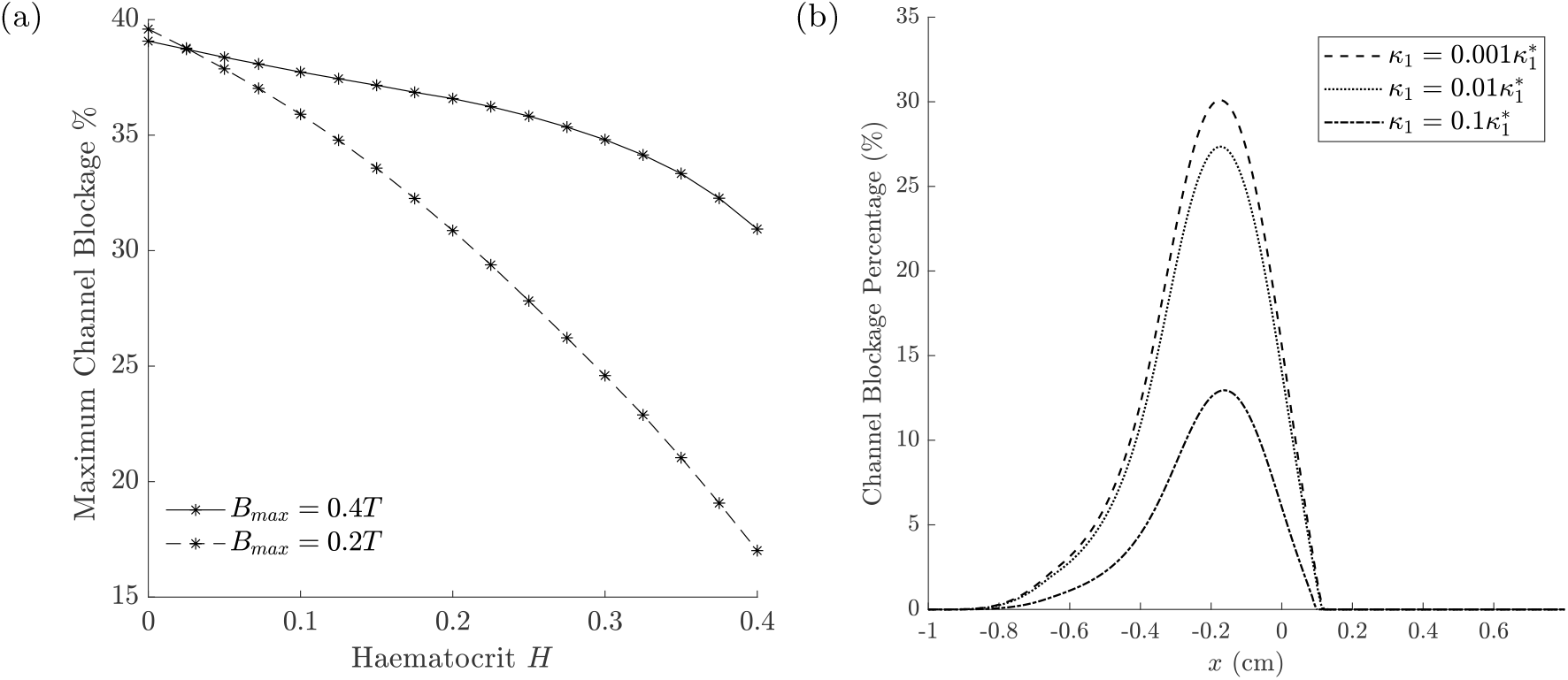
(a) Maximum aggregate height decreases from 40% as haematocrit increases. Parameters: *γ* = 0.1*γ*\* and no extravasation, all other parameters in Table 2. (b) *In vivo* delivery: aggregate interface profiles decrease for increasing height dependent extravasation parameter *κ*_1_. Parameter values: *H* = 0.2, *B_max_* = 0.4*T*, *γ* = 0 (no erosion) and at *t* = 50*t*\*, *t*\* = 0.11s all others as in Table 2.

### Model extension to *in vivo* delivery: extravasation allows a greater number of cells to be captured with less channel obstruction

Including extravasation through the vessel wall reduces the height of the aggregate at all axial positions, as illustrated in Figure 9b. To compare the effect of the two extravasation kinetics, constant and height dependent (defined in Eq. (4)), we plot the height of the aggregate across the channel over time, Fig. 10. To quantitatively compare we choose an extravasation rate such that *κ*_1_*h* is comparable to *κ*_2_. Constant extravasation leads to greater obstruction of the channel and the cells persist in the channel for a longer time and over a larger area. Height dependent extravasation leads to an aggregate which clears from the vessel more rapidly.

**Figure 10:**
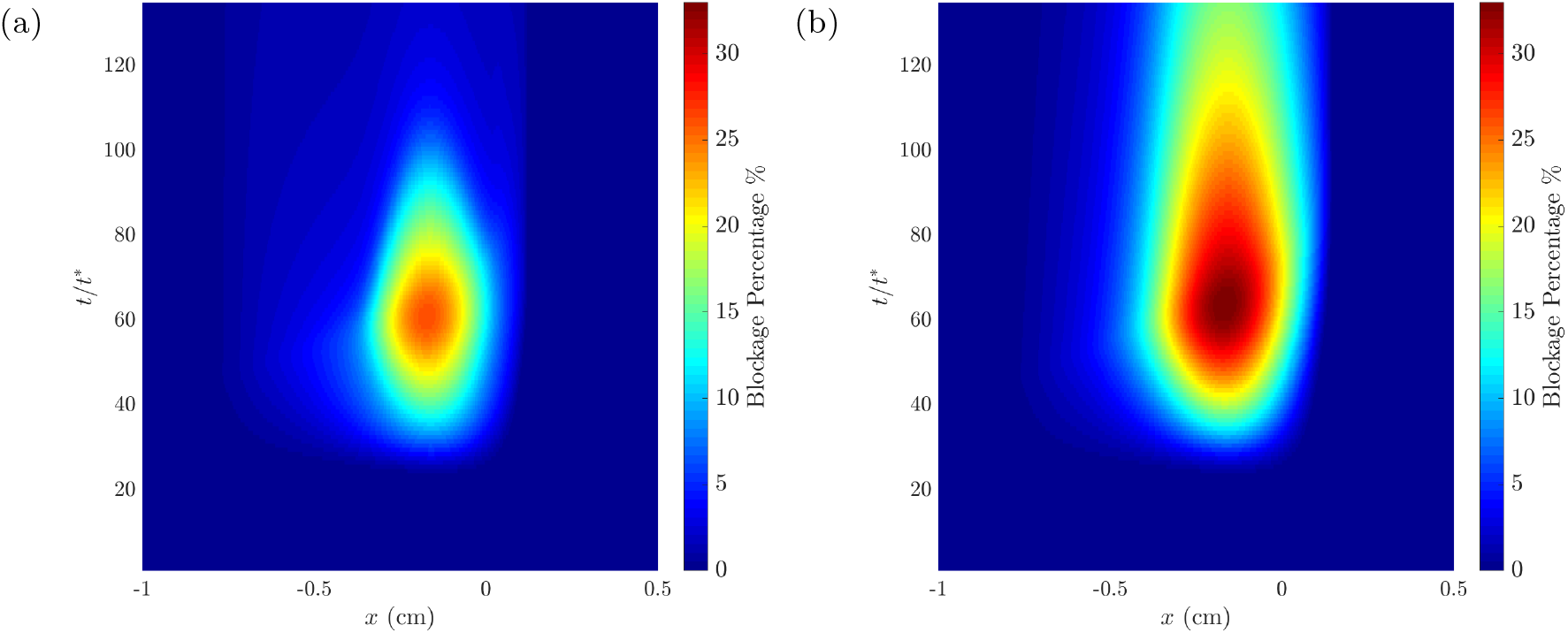
Channel blockage percentage across the centre of the channel over time. Left: height dependent extravasation 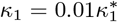, right: constant extravasation 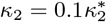. Constant extravasation leads to much greater channel blockage and the aggregate persisting in the channel for a longer time. Parameter values: *H* = 0.2, *B_max_* = 0.4*T, γ* = 0 and all others as in Table 2.

## 3 Discussion

The use of magnetic particles for medical applications is an expanding field with current and potential applications in areas such as imaging or cancer hyperthermia [17, 25] moving towards the clinic. Use of superparamagetic iron oxide nanoparticle (MNPs or SPIONs) are commonly integrated into MRI imaging as a contrast agent, with safety and efficacy in the clinical setting being consistently demonstrated [9, 35]. Novel injectable applications involve using labelled MNPs to tag to specific receptors on cells such as mechanosensitive ion channels or growth factors which can then be activated remotely for regenerative medicine or cell based therapies [27]. In many of these applications, targeting of the particles with and without cells to specific areas in the body is required and the use of magnetic fields to target injected cells to a specific region may become a feasible option [26].

In this study we have highlighted the potential of magnetic trapping by labelling the cells with MNPs and demonstrating their capture in a flow system using an external magnet. In addition, our data has highlighted the benefits of combined modelling approaches, e.g. experimental and theoretical, for optimising the conditions for magnetic targeting *in vivo* and *in vitro*. Although previous work has shown how MNPs can be trapped within an *in vitro* experimental model [8] these studies did not take into account the presence of blood cells in the vascular supply which may impact on the ultimate trapping and extravasation of the cells into the tissue. In this current study, we highlight how blood cells interfere with the cells accumulation in the region of the highest magnetic field with a clear relationship between the size of the magnetic field, number of blood cells and number of the MNPs. This experimental model has limitations as the cells become trapped onto the surface of the simulated vessel, whereas, *in vivo* these cells would move through this outer wall into the tissues at a given rate. The mathematical model presented allows us to consider these elements both in terms of aggregation and also extravasation. Although the trapping percentages are not yet optimised, particularly for higher blood concentrations, the improvements made in this system indicate the potential effectiveness, with key benefits for clinical introduction.

The *in vitro* experimental model considered a single vessel and analysed the key processes of magnetic targeting. However, *in vitro* experimental vessel models with more complex geometries could be employed to study how magnetic targeting would affect MSC delivery through a network of vessels [34] or to include extravasation [19] using co-culture endothelial wall mimics. This would allow choices such as magnet position and location of MSC injection to be examined. The use of diamagnetism in such a system, as opposed to paramagnetism, has also received interest; with the main difference being a repulsive force acting on the cells instead of attractive [10, 30]. Although this research tends to focus more on microfluidic arrays, as opposed to *in vitro* cell therapy targeting.

The labelling protocol utilised in this experiment has been shown in numerous publications to provide a labelling efficiency > 95% [1, 16, 27]; however, the homogeneity of labelling density is not yet standardised, as highlighted in the images obtained from Prussian blue staining. Ideally, the protocol could be adapted such that all cells are labelled above the threshold value mentioned in the results section. Our results show the effect of labelling concentrations on our ability to trap since the majority of trapped cells showed a higher labelling efficiency. One method of attempting this would be to label a greater number of cells, and isolate cells above a certain magnetic threshold providing samples of greater homogeneity. Similar concepts have been developed, whereby a system can be made which traps particles in a flow system using a pneumatically operated magnetic arm. Furthermore, finite time experiments would enable us to determine the optimum flow time to maximise trapping efficiency. The data suggest a short application time is required for maximum trapping to occur. This again is a key factor if considering development into clinical applications, as a short treatment injectable solution would be necessary for widespread adoption of this technique. Due to the ongoing improvements surrounding this trapping method, further work is still required to assess optimum labelling methods; and also to further understand the process by which the particles cross cell membranes. This is key to allowing incorporation of these methods into clinical applications.

The mathematical model used in this study extends continuum models of nanoparticle delivery [14, 33] to focus on the capture of cells. The arrival of MSCs at the magnetic source leads to obstruction of the channel through the growth of a cell aggregate. The solid aggregate grows due to MSC arrival and cells are removed from its surface by fluid shear stress. This simple mathematical model can reproduce the capture trends shown *in vitro*: fewer MSCs are captured when higher percentages of RBCs are present and that increasing magnetic field strength increases capture. The theoretical model, however, goes further and offers insight into predicting *in vivo* capture of MSCs through the inclusion of cell extravasation. Two simple extravasation functional forms were tested and demonstrated different qualitative capture of MSCs. Firstly for constant extravasation greater numbers of MSCs were captured in the channel over a larger region, whereas for height dependent extravasation cells exited the vessel faster and minimised blockage.

This mathematical model can offer insight into the key factors governing MSC delivery by making simplifying assumptions, which have limitations, and there are a number of potential extensions which could be made to the model. For example in our model we assumed unaggregated MSCs are dilute, therefore, neglecting their effect on the fluid flow. Previous models have included the effect which MNPs have on fluid flow [5, 13]. This was noted to have the potential to induce mixing in extreme cases [13]. Inclusion of this would add a magnetic force on the fluid flow, which could slow down the flow and change cell capture dynamics. Furthermore, we neglected magnetic interaction between internal MNPs and between neighbouring cells. Construction of a continuum representation for the effect of these interactions on a density field of superparamagnetic MNPs is a current open question.

The model further assumes that the aggregated cells are solid which neglects internal flow. The aggregate of MSCs at the magnet is likely to be porous rather than solid and RBCs may also be accidentally captured adding further heterogeneity to the mass. Porous heterogenous clot formation has been studied in the context of thrombosis, where interclot flow along with RBC entrapment were shown to be essential in understanding clot stability and possible fracture [7, 39]. Examining the internal aggregate structure and tracking the RBC distribution explicitly would be a valuable next step for the model.

The mathematical model enables us to extend and predict from the parameters obtained from our experimental model, however, validation with more complex experimental models would be useful to confirm our predictions for aggregate growth, fluid erosion and extravasation rates. Some examples of *in vitro* models have been developed [4, 19] which quantify and visualise extravasation of other cell types or the effect of the magnetic force on extravasation speed [29]. These models could be used to parameterise the mathematical model. Similarly, the *in vitro* model discussed in this paper could be employed to determine the MSCs susceptibility to erosion by varying flow rates over pre-captured cells. Quantification of these parameters would allow the model to provide a prediction of aggregation dynamics, providing insight on dosage levels required to ensure that MSC aggregation is kept within safe limits while maintaining therapeutic efficacy.

## 4 Conclusion

A combined experimental and mathematical approach to modelling has been presented. We have demonstrated the role of circulating blood cells and magnetic field strength on MSC capture and vessel obstruction. The mathematical model is able to replicate trends of *in vitro* experimental data and predict the interplay between the key mechanisms of delivery. It is able to provide an upper bound on vessel obstruction in the *in vitro* scenario and to bridge between *in vitro* and *in vivo* delivery through the inclusion of extravasation. We have demonstrated that extravasation always reduces the level of MSC aggregation and is key to cell clearance from the vessel. This highlights directions for new experimental investigation and potential use of magnetic targeting for optimising cell delivery systems for regenerative medicine and other applications.

## Data Availability

Raw experimental data is provided in the supplementary material. COMSOL and MATLAB files for figure production can be found in the repository https://github.com/Edwina-Yeo/Magnetic-Cell-Delivery-Simulations.

## Authors’ Contributions

Conceived the project: SLW & AEH; performed small magnet experiments: AJS; performed large magnet experiment: ATS; supervised ATS & wrote relevant experimental sections: HM; constructed the mathematical model: EY, JMO & SLW; performed simulations and wrote remainder of paper EY; provided comments on manuscript: SLW, HM & AEH. All authors gave final approval for publication and agree to be held accountable for the work performed therein.

## Appendices

## A Full Governing Equations

The full mathematical model is described below.

## Fluid Flow

The fluid is assumed to be slow such that inertia can be neglected and Stokes and continuity equations govern the flow

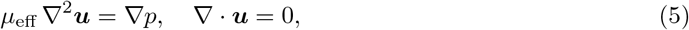

where ***u*** and *p* are the fluid velocity and pressure respectively and *μ*_eff_ is an effective viscosity, defined below.

## Effective viscosity

The viscosity is evaluated relative to the viscosity of water *μ*_water_ and is a function of RBC fraction, we use the empirical fitting [31], as follows

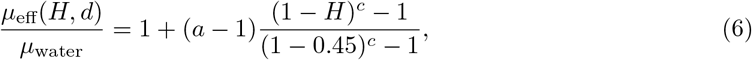

where coefficients *a* and *c* are functions of vessel radius. Throughout this model we use the values of *a* = 3.19 and *c* = −0.8 (for the pipe radius used *in vitro*) such that the viscosity is a function of *H* alone. Although the range of experimental data compiled in Pries’s papers is for diameters from 10*μm* to 1000*μm* and the *in vitro* pipe has diameter 1650*μm* we can use this fitting since, as stated in Pries (1996) [31], the viscosity does not change significantly beyond this value.

## Stem cell transport

Stem cell concentration *c* in the channel is governed by and advection diffusion equation as in [14]

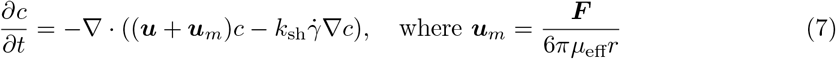

where *D*_sh_ is the shear induced diffusion coefficient, ***u***_*m*_ is the magnetic induced velocity, ***F*** is the magnetic force on a cell radius *r* tagged with *n* NPs. The diffusion coefficient is defined *D_sh_* = *kr*^2^*H*^2^, where *r* is the stem cell radius [23], *k* is a constant and 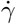 is the local shear rate defined as the 2-norm of the strain rate tensor

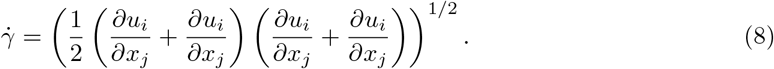

## Magnetic Field

The applied magnetic field is generated by an infinite cylindrical magnet of radius *R_mag_* and with magnetisation in the *y*-direction. The magnet is located directly below the channel, as *in vitro*, such that its centre is at *x* = 0, *y* = −*d* − *R_mag_*. This allows the magnetic field to be defined as

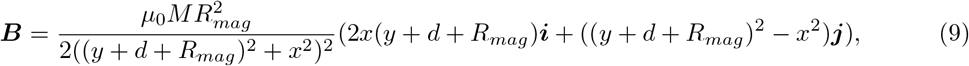

where *M* is the magnetisation of the external magnet and *μ*_0_ is the magnetic permeability of a vacuum. The magnetic moment of a cell is assumed to be saturated and at linear multiple of the moment of a NP ***m*** = *nm*_*NP*_ with the moment of a nanoparticle as 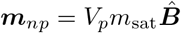 [33]. The force on a magnetic nanoparticle is obtained by approximating the nanoparticle as pure magnetic dipole [15]

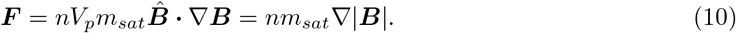

The latter equality is obtained through application of Ampèhre’s law, since there is no current. Hence using Eq. (10) the force on a cell can be expressed analytically [11], *viz*

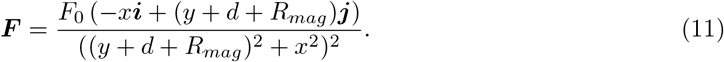

In the experimental model, the external field was provided by rectangular and cylindrical magnets, however, the theoretical magnet geometry was an infinite cylinder oriented perpendicular to the channel. The geometry difference is minimised by ensuring the radius of the theoretical magnet is similar to the experimental magnets.

## Aggregate interface velocities

The normal and tangential components of the aggregate interface velocity are defined by

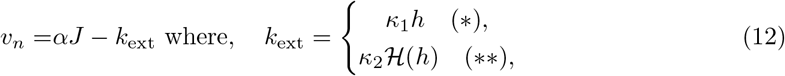

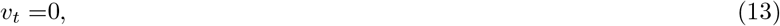

where *J* is the flux of stem cells relative to the moving boundary, *α* accounts for the conversion of cell concentration into solid aggregate, *k*_ext_ is the extravasation function, where cases of height dependent extravasation and constant extravasation are defined by (∗) and (∗∗) respectively.

All boundary conditions and initial conditions are summarised in Figure 11 and 12.

**Figure 11:**
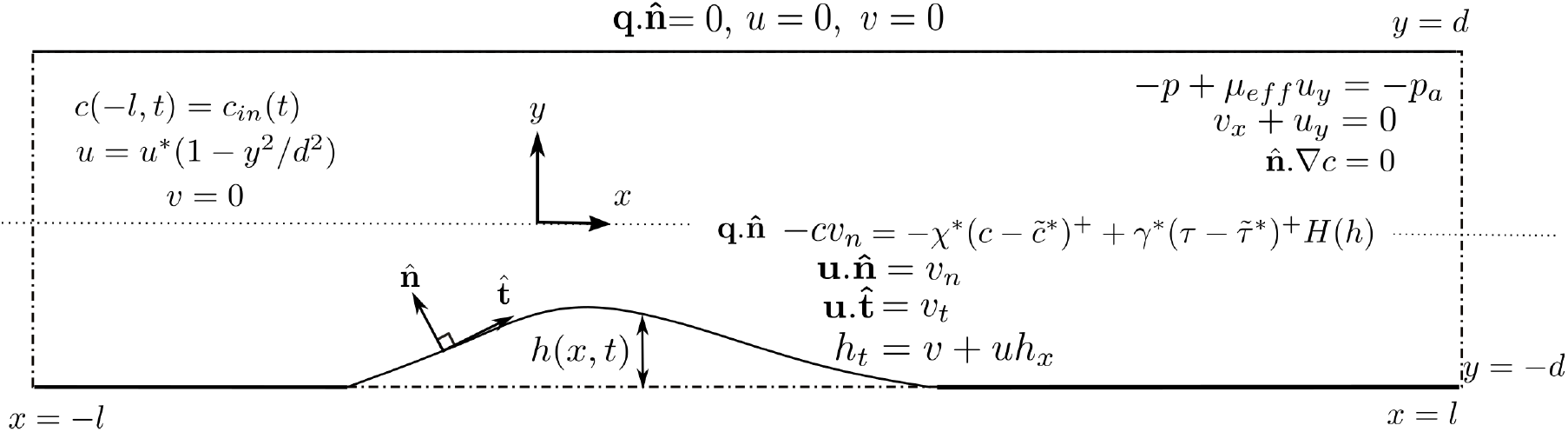
Boundary conditions on the system, where 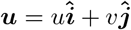

**Figure 12:**
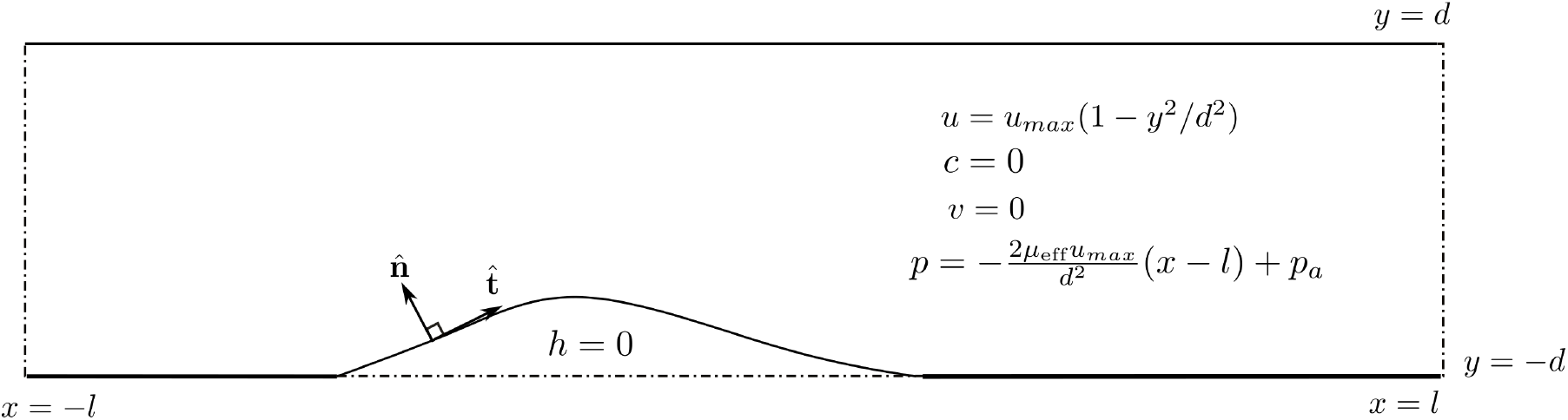
Initial conditions on the system

## A.1 Dimensionless Model

All simulations were carried out on the dimensionless model presented below. The results were presented in dimensional form for ease of comparison to experimental data. The dimensional model is obtained from Eqs. (5) - (12) by inserting the following scalings

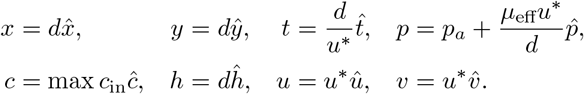

We take the cell concentration scale to be max *c*_in_ such that the largest inlet value is unity.

Inserting these scalings and dropping the hats we arrive at the dimensionless system

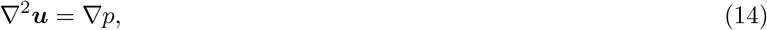

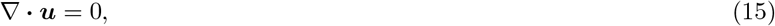

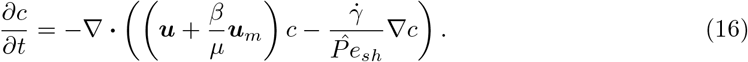

All dimensionless parameters and values used in simulations are defined in Tables 1 and 2. We note that we have left *μ* = *μ*_eff_*/μ*_water_ explicit in the model so that we can easily vary both the magnetic field strength and the RBC percentage. The dimensionless shear rate is

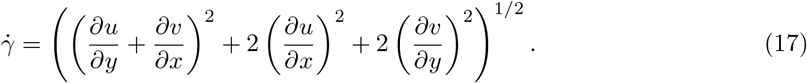

The dimensionless aggregate interface velocities are defined by

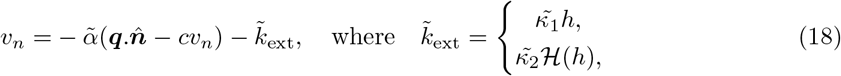

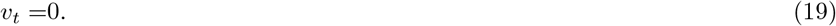

Dimensionless boundary conditions are

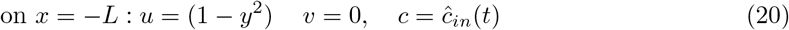

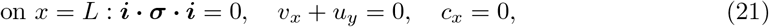

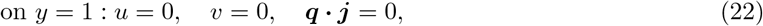

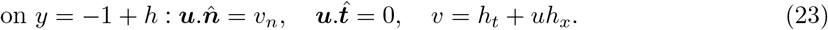

The flux condition on cell equation is

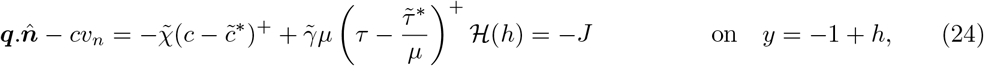

the dimensionless inlet condition 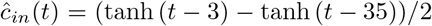. The dimensionless mag-netic velocity is

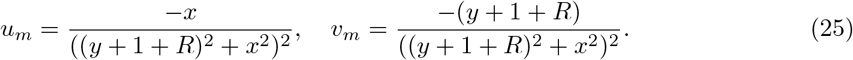

**Table 3:** Dimensionless parameter definitions and values used in simulations. The † marks the deviation from definition, the inverse Peclet number had to be increased to ensure numerical stability

| Parameter | Definition | Definition | Value |
| --- | --- | --- | --- |
| $1/\hat{P}e_{sh}$ | inverse shear Peclet number | $0.5H^2r_{RBC}^2/d^2$ | 0.1(†) |
| $\mu$ | relative increase in viscosity with Haematocrit | $\mu_{\text{eff}}/\mu_{\text{water}}$ | Defined in Eq (6) (Appendix) |
| $\beta$ | ratio of magnetic to fluid advection | $n\mu_0m_{sat}MR_{mag}^2/(6\pi\mu_{\text{water}}r d^3U)$ | 52, 105, 157, 210 |
| $\tilde{\alpha}$ | dimensionless cell flux conversion factor | $\alpha \max c_{\text{in}}$ | 0.1 |
| $\tilde{\kappa}_1$ | dimensionless extravasation rate | $\kappa_1 d/U$ | 0.001, 0.01, 0.1 |
| $\tilde{\kappa}_2$ | dimensionless constant extravasation rate | $\kappa_2/U$ | 0.1 |
| $\tilde{\chi}$ | dimensionless uptake rate | $\chi/U$ | 0.1 |
| $\tilde{\gamma}$ | dimensionless erosion factor | $\gamma\mu_{\text{water}}/d \max c_{\text{in}}$ | 0.1, 0.4, 0.8 |
| $\tilde{c}^*$ | Ratio of magnetic to fluid advection | $c^*/\max c_{\text{in}}$ | 1.1 |
| $\tilde{\tau}^*$ | dimensionless erosion threshold | $\tau^* d/\mu_{\text{water}}U$ | 4 |
| $R$ | ratio of magnet radius to channel half width | $R_{mag}/d$ | 4.24 |
| $L$ | ratio of channel half width to length | $l/d$ | 20 |

## B Numerical Verification

## B.1 Regularised Heaviside Functions

The flux condtion for the cells, Eq. (3), requires three Heaviside functions. Defining the numerical approximation to the Heaviside function by 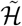 this is implemented as follows

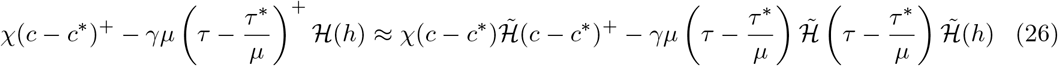

We use the inbuilt function ‘flc1hs(*x, δ*)’ in COMSOL [6] which has continuous first derivative and is defined as a fifth order polynomial. The function is equal to zero for *x* < −*δ* and one for *x > δ*. This is illustrated in Fig. 13 for varying *δ.* Since we need to avoid the approximate Heaviside function having postive values for any *x* < 0, as this would allow effects such as erosion to occur when there is no aggregate, we chose to use the Heaviside function shifted by *δ* ‘flc1hs(*x* − *δ, δ*)’ which is equal to zero for *x* < 0 and one for *x* > 2*δ*.

**Figure 13:**
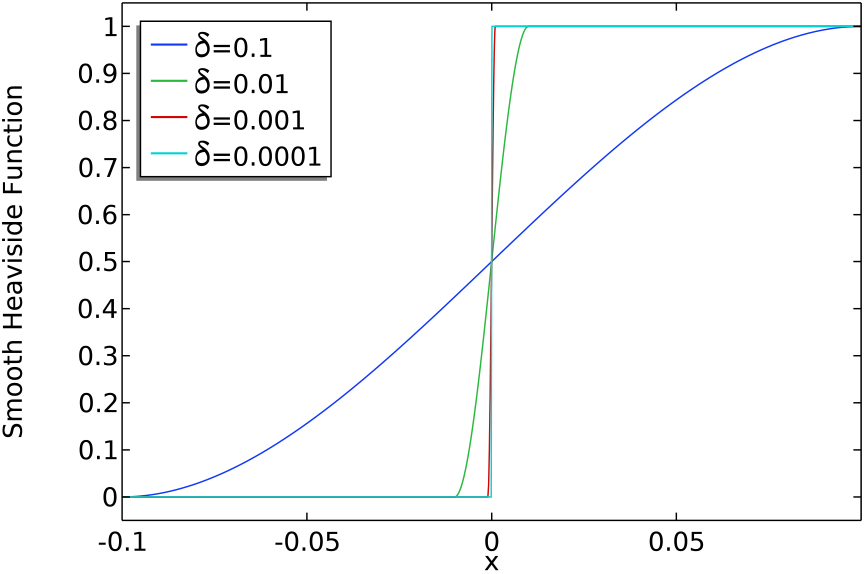
Plot of the regularised heaviside function ‘flc1hs(*x, δ*)’ for decreasing *δ*, as *δ* → 0 the regularised function tends to the analytic Heaviside function.

We have quantified the effect of reducing the smoothing parameter *δ* on the solution. As shown in Table 4 the change in the aggregate height attained is < 10^−3^. Hence we chose to use the *δ* = 0.1 as reducing only serves to increase the computational cost but no longer changes the solution significantly.

**Table 4:** Maximum difference in *h* is measured compared to the case *δ* = 10^−4^ as the smoothing parameter is reduced. Since the effect minimal on the height *h* we use the value *δ* = 10^−3^.

| $\delta$ | Maximum % change in $h$<br>relative to $\delta = 10^{-4}$ |
| --- | --- |
| 0.1 | 0.22 |
| 0.01 | $10^{-3}$ |
| 0.001 | $< 10^{-4}$ |

## B.2 Removing Shear induced Diffusion

In order to have tractable numerical simulations the cell advection - diffusion equation (16) was approximated as follows

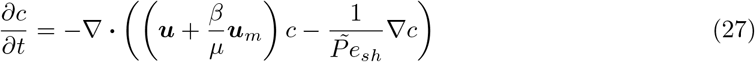

This removes the dependence on fluid shear and increases the inverse Peclet number, hence increasing the effect of diffusion. We choose 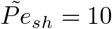.

## B.3 Mesh Convergence Tests

We construct a mesh with finer elements near the boundary then reduce the total number of elements while maintaining this structure. We quantify the errors as follows for a mesh with *n* elements

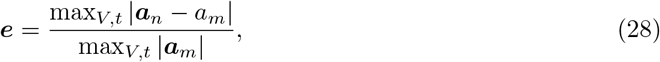

where ***a*** = (*c, u, v, p*) is a vector of the dependent variables. The fine comparison mesh has *m* = 13758 elements. We use the third mesh which has *n* = 4202 elements as the percentage error is ≤ 1.59%(as shown in Figures 14, 15 and 16). We note that we restrict our study to values of 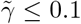 since the coupling of the fluid shear to the moving boundary makes the problem stiff, hence we restrict gamma to ensure reasonable convergence (poor convergence shown in Fig 17 for 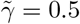). A section of the mesh is shown in Figure 18.

**Figure 14:**
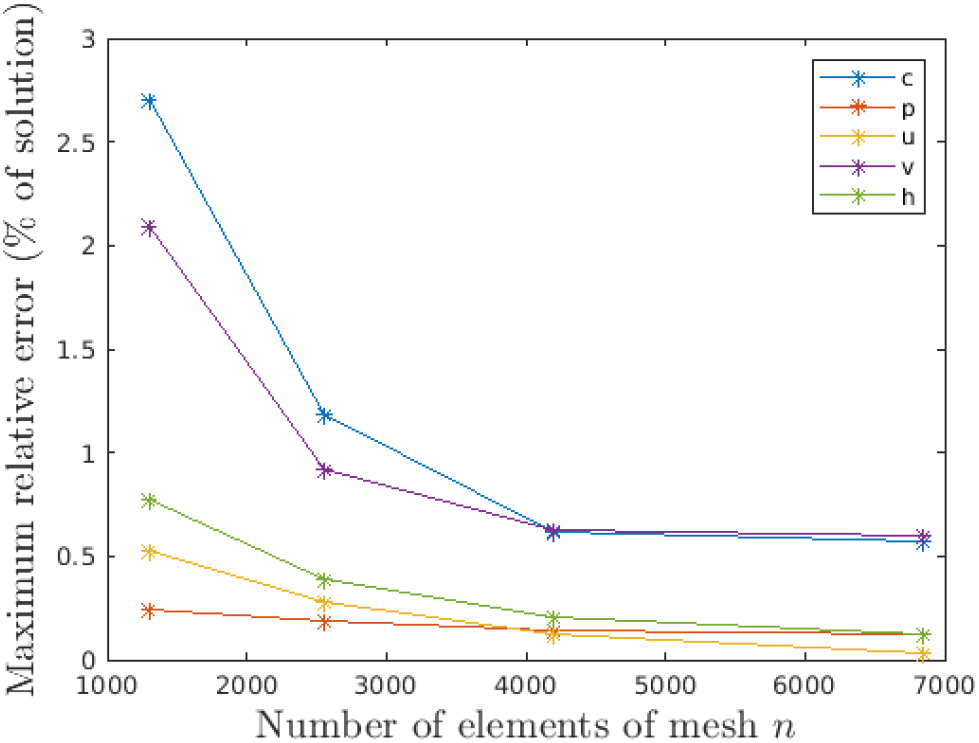
Percentage error of each variable relative to the maximum value of that variable. Parameters: *β* = 0.4*T*, 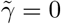, 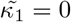. The third mesh provides < 1% change in all variables.

**Figure 15:**
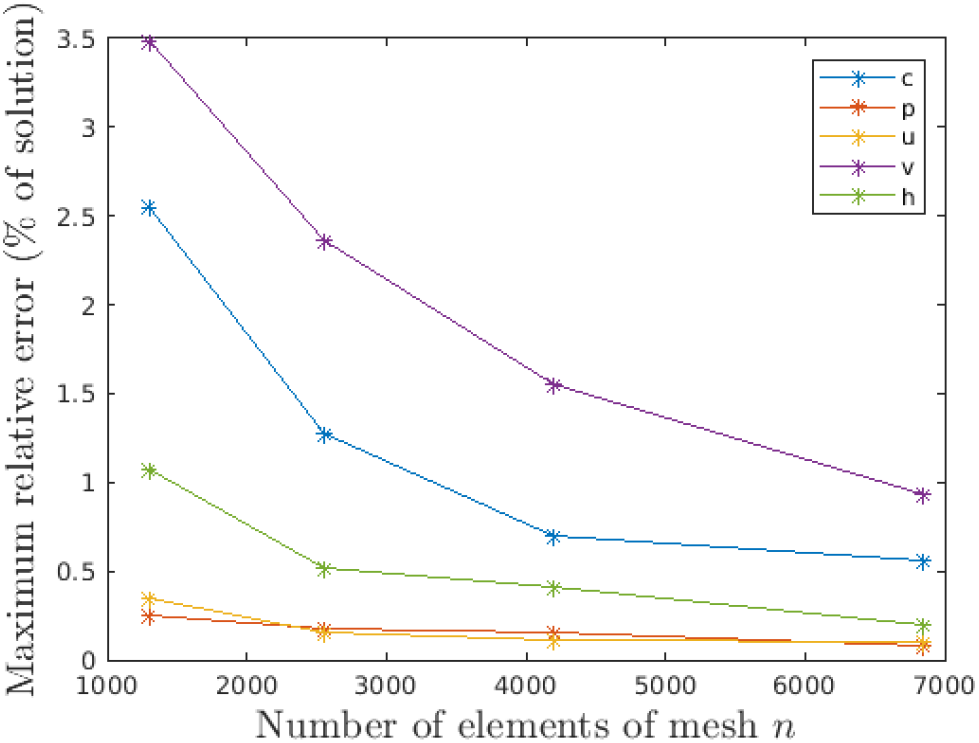
Percentage error of each variable relative to the maximum value of that variable. Parameters: *β* = 0.4*T*, 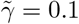, 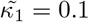. The third mesh provides all variables error ≤ 1.5%.

**Figure 16:**
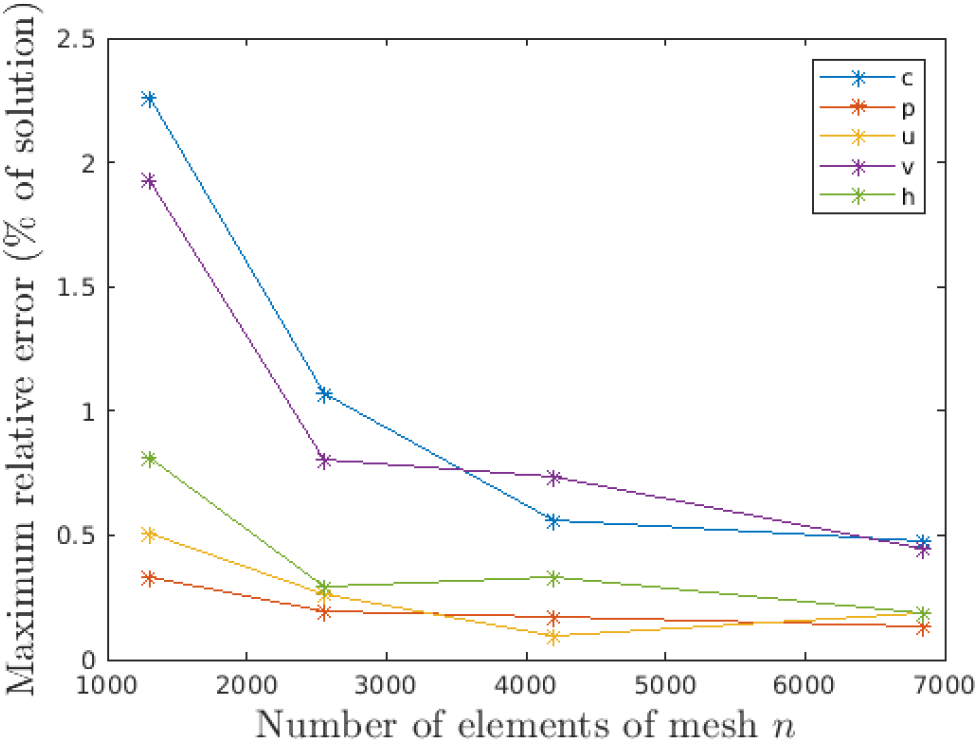
Percentage error of each variable relative to the maximum value of that variable. Parameters: *β* = 0.4*T*, 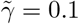, 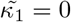. The third mesh provides all variables error < 1%.

**Figure 17:**
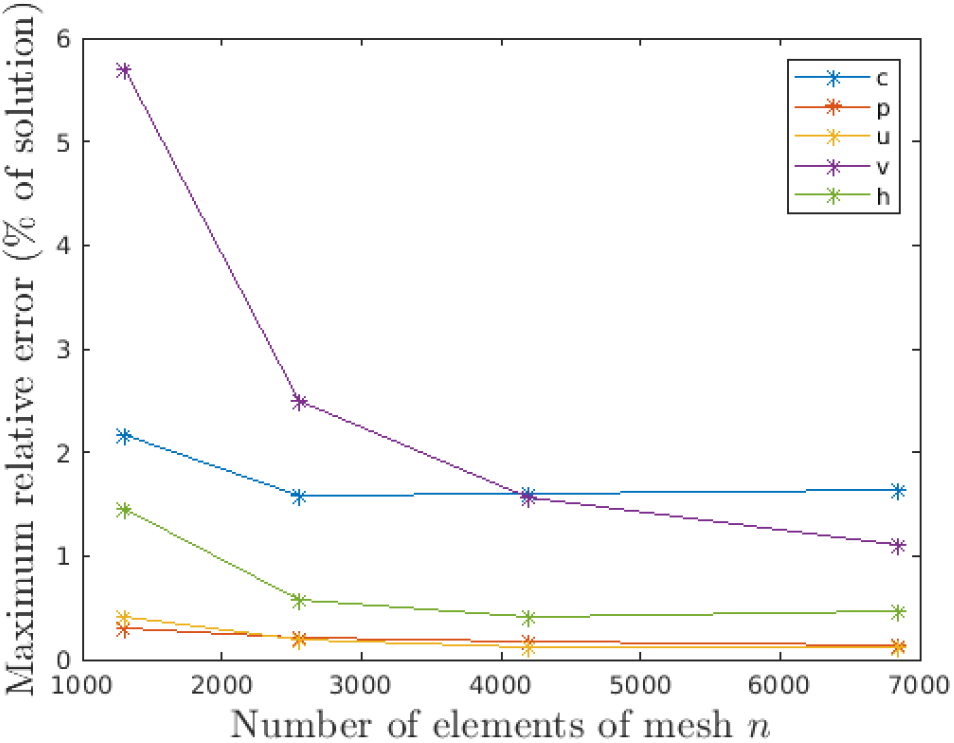
Percentage error of each variable relative to the maximum value of that variable. Parameters: *β* = 0.4*T*, 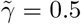, 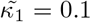. The solution does not have good convergence properties as 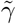 is larger, we restrict the value of 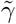 considered to achieve better stability properties.

**Figure 18:**
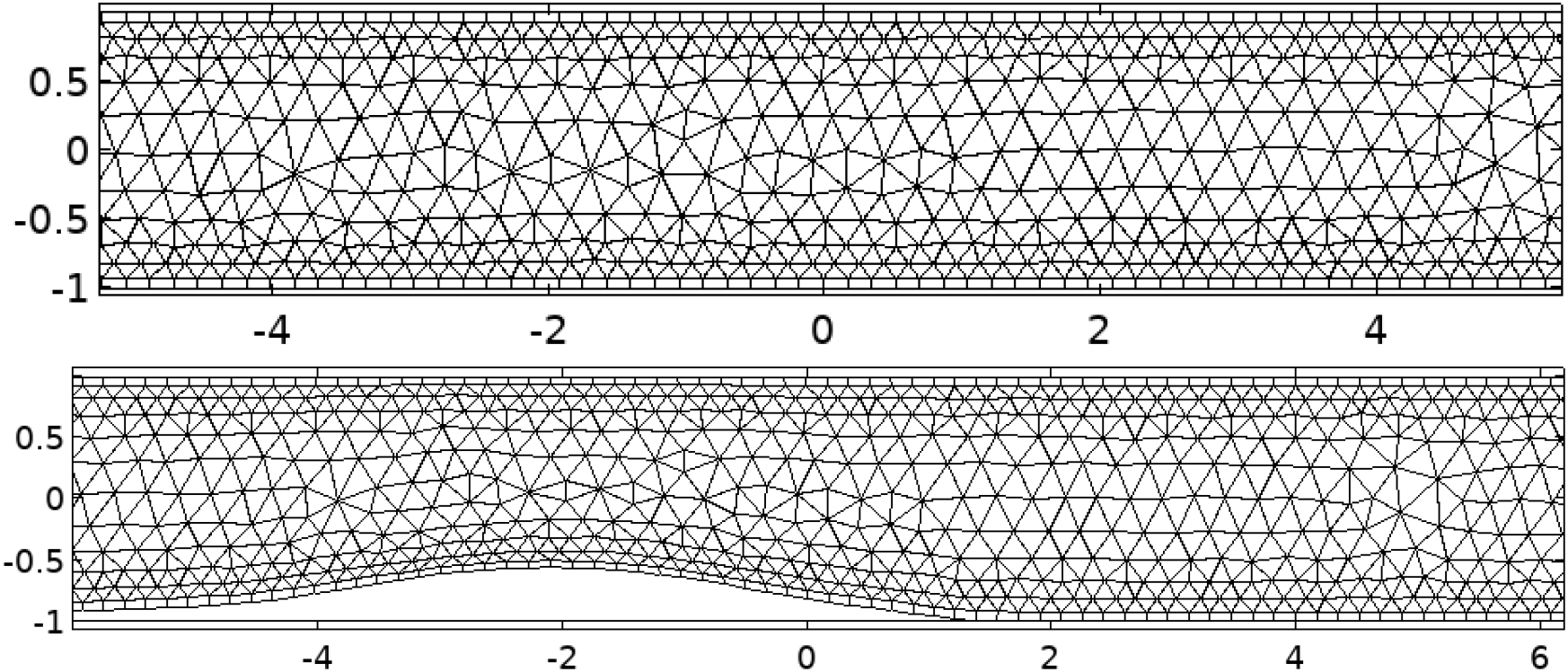
Examples of the mesh used to compute. At *t* = 0 and after deformation at *t* = 50*t*\*

