## Supplementary material for "Experimental and mathematical modelling of magnetically labelled mesenchymal stromal cell delivery": Table S2

| RAW DATA | Trapped | | | | | Untrapped | | | | |
|---|---|---|---|---|---|---|---|---|---|---|
| Group | Sample 1 | Sample 2 | Sample 2 | Average | Standard Deviation | Sample 1 | Sample 2 | Sample 2 | Average | Standard Deviation |
| 0% RBC - MNPs | 0.4 | 0.4 | 0.5 | 0.433333333 | 0.057735027 | 30.4 | 25.8 | 23.8 | 26.66666667 | 3.384277372 |
| 0% RBC + MNP | 150 | 169 | 238 | 185.6666667 | 46.30694692 | 180 | 204 | 87 | 157 | 61.79805822 |
| 1% RBC - MNPs | 0.8 | 0.9 | 1.2 | 0.966666667 | 0.2081666 | 26.2 | 22.5 | 24.5 | 24.4 | 1.852025918 |
| 1% RBC + MNP | 64 | 43 | 114 | 73.66666667 | 36.47373484 | 55 | 41 | 37 | 44.33333333 | 9.451631253 |
| 5% RBC - MNPs | 0.9 | 0.7 | 0.8 | 0.8 | 0.1 | 18.9 | 24.6 | 18.7 | 20.73333333 | 3.350124376 |
| 5% RBC + MNP | 49 | 20 | 28 | 32.33333333 | 14.97776129 | 88 | 72 | 56 | 72 | 16 |
| 10% RBC - MNPs | 1.5 | 1.3 | 2 | 1.6 | 0.360555128 | 27.3 | 18.4 | 15 | 20.23333333 | 6.351640208 |
| 10% RBC + MNP | 53 | 31 | 45 | 43 | 11.13552873 | 68 | 98 | 50 | 72 | 24.24871131 |
| 20% RBC - MNP | 1.3 | 3 | 2.9 | 2.4 | 0.953939201 | 29.5 | 32.7 | 17.1 | 26.43333333 | 8.239741096 |
| 20% RBC + MNPs | 8 | 24 | 33 | 21.66666667 | 12.66227994 | 57 | 51 | 63 | 57 | 6 |
| 40% RBC + MNP | 27 | 21 | 22 | 23.33333333 | 3.214550254 | 105 | 59 | 62 | 75.33333333 | 25.73583753 |
| 40% RBC - MNP | 1.3 | 1.9 | 2.7 | 1.966666667 | 0.702376917 | 18.7 | 15.8 | 10 | 14.83333333 | 4.429823172 |

| PROCESSED DATA | | | | |
|---|---|---|---|---|
| Group | Trapped | Untrapped | Percentage Trapp | Percentage Untrapped |
| 0% RBC - MNPs | 0.4333333 | 26.6666667 | 1.60 | 98.40 |
| 0% RBC + MNP | 185.66667 | 157 | 54.18 | 45.82 |
| 1% RBC - MNPs | 0.9666667 | 24.4 | 3.81 | 96.19 |
| 1% RBC + MNP | 73.666667 | 44.3333333 | 62.43 | 37.57 |
| 5% RBC - MNPs | 0.8 | 20.7333333 | 3.72 | 96.28 |
| 5% RBC + MNP | 32.333333 | 72 | 30.99 | 69.01 |
| 10% RBC - MNPs | 1.6 | 20.2333333 | 7.33 | 92.67 |
| 10% RBC + MNP | 43 | 72 | 37.39 | 62.61 |
| 20% RBC - MNP | 2.4 | 26.4333333 | 8.32 | 91.68 |
| 20% RBC + MNPs | 21.666667 | 57 | 27.54 | 72.46 |
| 40% RBC + MNP | 23.333333 | 75.3333333 | 23.65 | 76.35 |

| STDEV | Trapped | | | | | Untrapped | | | | |
|---|---|---|---|---|---|---|---|---|---|---|
| Group | Sample 1 | Sample 2 | Sample 2 | Average | STDEV | Sample 1 | Sample 2 | Sample 2 | Average | STDEV |
| 0% RBC - MNPs | 1.2987013 | 1.52671756 | 2.057613169 | 1.627677342 | 0.389398851 | 98.7012987 | 98.4732824 | 97.9423868 | 98.37232266 | 0.389398851 |
| 0% RBC + MNP | 45.454545 | 45.308311 | 73.23076923 | 54.66454189 | 16.07899077 | 54.5454545 | 54.691689 | 26.7692308 | 45.33545811 | 16.07899077 |
| 1% RBC - MNPs | 2.962963 | 3.84615385 | 4.6692607 | 3.826125837 | 0.853325163 | 97.037037 | 96.1538462 | 95.3307393 | 96.17387416 | 0.853325163 |
| 1% RBC + MNP | 53.781513 | 51.1904762 | 75.49668874 | 60.15622585 | 13.34824778 | 46.2184874 | 48.8095238 | 24.5033113 | 39.84377415 | 13.34824778 |
| 5% RBC - MNPs | 4.5454545 | 2.76679842 | 4.102564103 | 3.804939022 | 0.92592652 | 95.4545455 | 97.2332016 | 95.8974359 | 96.19506098 | 0.92592652 |
| 5% RBC + MNP | 35.766423 | 21.7391304 | 33.33333333 | 30.27962904 | 7.495670024 | 64.2335766 | 78.2608696 | 66.6666667 | 69.72037096 | 7.495670024 |
| 10% RBC - MNPs | 5.2083333 | 6.59898477 | 11.76470588 | 7.857341329 | 3.454576844 | 94.7916667 | 93.4010152 | 88.2352941 | 92.14265867 | 3.454576844 |
| 10% RBC + MNP | 43.801653 | 24.0310078 | 47.36842105 | 38.40036057 | 12.57136365 | 56.1983471 | 75.9689922 | 52.6315789 | 61.59963943 | 12.57136365 |
| 20% RBC - MNP | 4.2207792 | 8.40336134 | 14.5 | 9.041380188 | 5.16922586 | 95.7792208 | 91.5966387 | 85.5 | 90.95861981 | 5.16922586 |
| 20% RBC + MNPs | 12.307692 | 32 | 34.375 | 26.2275641 | 12.11331001 | 87.6923077 | 68 | 65.625 | 73.7724359 | 12.11331001 |
| 40% RBC + MNP | 20.454545 | 26.25 | 26.19047619 | 24.29834055 | 3.328957241 | 79.5454545 | 73.75 | 73.8095238 | 75.70165945 | 3.328957241 |
| 40% RBC - MNP | 6.5 | 10.7344633 | 21.25984252 | 12.83143527 | 7.600079371 | 93.5 | 89.2655367 | 78.7401575 | 87.16856473 | 7.600079371 |

Table S2: Raw data used to calculate the trapping results for the large magnet (strength 0.4T), Figure 7b.
