## Supplementary material for "Experimental and mathematical modelling of magnetically labelled mesenchymal stromal cell delivery": Table S1

| RAW DATA | Trapped | | | | | Untrapped | | | | |
|---|---|---|---|---|---|---|---|---|---|---|
| Group | Sample 1 | Sample 2 | Sample 2 | Average | Standard Deviation | Sample 1 | Sample 2 | Sample 2 | Average | Standard Deviation |
| 0% RBC - MNPs | 17 | 25 | 30 | 24 | 6.557438524 | 96 | 138 | 116 | 116.66667 | 21.00793501 |
| 0% RBC + MNP | 133 | 87 | 104 | 108 | 23.2594067 | 95 | 128 | 86 | 103 | 22.11334439 |
| 1% RBC - MNPs | 9 | 11 | 22 | 14 | 7 | 2110 | 1380 | 1700 | 1730 | 365.9234893 |
| 1% RBC + MNP | 141 | 127 | 174 | 147.3333333 | 24.13158373 | 740 | 930 | 780 | 816.66667 | 100.166528 |
| 5% RBC - MNPs | 5 | 6 | 8 | 6.333333333 | 1.527525232 | 1150 | 1140 | 1370 | 1220 | 130 |
| 5% RBC + MNP | 125 | 75 | 101 | 100.3333333 | 25.00666578 | 1010 | 1030 | 910 | 983.33333 | 64.29100507 |
| 10% RBC - MNPs | 11 | 13 | 8 | 10.66666667 | 2.516611478 | 560 | 1030 | 840 | 810 | 236.4318084 |
| 10% RBC + MNP | 85 | 25 | 58 | 56 | 30.0499584 | 1080 | 1860 | 1170 | 1370 | 426.7317659 |
| 20% RBC - MNP | 29 | 46 | 33 | 36 | 8.888194417 | 5880 | 6530 | 5550 | 5986.6667 | 498.6314604 |
| 20% RBC + MNPs | 108 | 56 | 42 | 68.66666667 | 34.77547028 | 1730 | 700 | 460 | 963.33333 | 674.7098142 |
| 40% RBC + MNP | 56 | 43 | 31 | 43.33333333 | 12.50333289 | 2180 | 3170 | 2910 | 2753.3333 | 513.2575702 |

| PROCESSED DATA | | | | |
|---|---|---|---|---|
| Group | Trapped | Untrapped | Percentage Trapped | Percentage Untrapped |
| 0% RBC - MNPs | 24 | 116.6667 | 17.06 | 82.94 |
| 0% RBC + MNP | 108 | 103 | 51.18 | 48.82 |
| 1% RBC - MNPs | 14 | 1730 | 0.80 | 99.20 |
| 1% RBC + MNP | 147.33333 | 816.6667 | 15.28 | 84.72 |
| 5% RBC - MNPs | 6.3333333 | 1220 | 0.52 | 99.48 |
| 5% RBC + MNP | 100.33333 | 983.3333 | 9.26 | 90.74 |
| 10% RBC - MNPs | 10.666667 | 810 | 1.30 | 98.70 |
| 10% RBC + MNP | 56 | 1370 | 3.93 | 96.07 |
| 20% RBC - MNP | 36 | 5986.667 | 0.60 | 99.40 |
| 20% RBC + MNPs | 68.666667 | 963.3333 | 6.65 | 93.35 |
| 40% RBC + MNP | 43.333333 | 2753.333 | 1.55 | 98.45 |

| STDEV | Trapped | | | | | Untrapped | | | | |
|---|---|---|---|---|---|---|---|---|---|---|
| Group | Sample 1 | Sample 2 | Sample 2 | Average | STDEV | Sample 1 | Sample 2 | Sample 2 | Average | STDEV |
| 0% RBC - MNPs | 15.044248 | 15.33742 | 20.54794521 | 16.97653877 | 3.096400479 | 84.95575221 | 84.66257669 | 79.45205479 | 83.023461 | 3.096400479 |
| 0% RBC + MNP | 58.333333 | 40.46512 | 54.73684211 | 51.17843057 | 9.450662128 | 41.66666667 | 59.53488372 | 45.26315789 | 48.821569 | 9.450662128 |
| 1% RBC - MNPs | 0.4247286 | 0.790798 | 1.277584204 | 0.831036946 | 0.427849309 | 99.57527135 | 99.20920201 | 98.7224158 | 99.168963 | 0.427849309 |
| 1% RBC + MNP | 16.00454 | 12.01514 | 18.23899371 | 15.41955706 | 3.152895743 | 83.9954597 | 87.98486282 | 81.76100629 | 84.580443 | 3.152895743 |
| 5% RBC - MNPs | 0.4329004 | 0.52356 | 0.580551524 | 0.512337389 | 0.074462574 | 99.56709957 | 99.47643979 | 99.41944848 | 99.487663 | 0.074462574 |
| 5% RBC + MNP | 11.013216 | 6.78733 | 9.990108803 | 9.26355166 | 2.204640813 | 88.98678414 | 93.21266968 | 90.0098912 | 90.736448 | 2.204640813 |
| 10% RBC - MNPs | 1.9264448 | 1.246405 | 0.943396226 | 1.372081887 | 0.503430458 | 98.07355517 | 98.7535954 | 99.05660377 | 98.627918 | 0.503430458 |
| 10% RBC + MNP | 7.2961373 | 1.32626 | 4.723127036 | 4.448508107 | 2.994398218 | 92.70386266 | 98.67374005 | 95.27687296 | 95.551492 | 2.994398218 |
| 20% RBC - MNP | 0.4907768 | 0.699513 | 0.591080064 | 0.593790076 | 0.104394685 | 99.50922322 | 99.30048662 | 99.40891994 | 99.40621 | 0.104394685 |
| 20% RBC + MNPs | 5.8759521 | 7.407407 | 8.366533865 | 7.216631131 | 1.256203027 | 94.12404788 | 92.59259259 | 91.63346614 | 92.783369 | 1.256203027 |
| 40% RBC + MNP | 2.5044723 | 1.338313 | 1.054063244 | 1.632282873 | 0.768593037 | 97.49552773 | 98.6616869 | 98.94593676 | 98.367717 | 0.768593037 |

Table S1: Raw data used to calculate the trapping results for the small magnet (strength 0.2T), Figure 7a.
